# N-acetyltransferase 10 promotes mRNA stability of immune response factors to modulate Zika virus infection

**DOI:** 10.1101/2025.07.17.665358

**Authors:** Shawn Gianola, Nicholas Lane, Kristen Kaytes, Cara T. Pager

## Abstract

Zika virus (ZIKV) is a re-emerging mosquito-borne flavivirus that poses serious risks to human health. In previous work, we used RNA immunoprecipitation and mass spectrometry to show that more than 30 distinct RNA modifications, or chemical moieties, were present on the RNA genome of ZIKV. Among these, N4-acetylcytosine (ac^4^C) was one of the most abundant modifications. In this study, we investigated the role of N-acetyltransferase 10 (NAT10), the writer enzyme that acetylates cytidine, in ZIKV gene expression. Using NAT10 knockout cell lines, RNA interference (RNAi), and overexpression rescue strategies, we found that loss of NAT10 led to increased levels of ZIKV protein and RNA. However, the production of infectious virus particles was not significantly affected. Interestingly, in NAT10-deficient cells compared to wild-type (WT) cells, ZIKV protein and RNA were detectable earlier during infection, suggesting that the loss of NAT10 facilitated increased viral replication. Despite this increase, ZIKV RNA was more rapidly degraded, although the accumulation of small flaviviral RNAs was not significantly altered by the absence of NAT10. Further analysis of key components of the innate immune response revealed that, in the absence of NAT10 and during early infection, *STAT1*, *IFIT1*, and *MX1* mRNA transcripts were rapidly degraded, leading to reduced expression of the respective innate immune proteins. Taken together, our findings demonstrate that NAT10, the ac^4^C writer enzyme, modulates the stability of specific innate immune mRNAs and thereby plays a regulatory role in ZIKV infection dynamics.

**IMPORTANCE:** RNA modifications are chemical groups that are deposited post-transcriptionally on RNA. In recent years, RNA modifications in viral RNAs have been shown to have profound effects on viral gene expression and hence viral function. Indeed, numerous studies have investigated the role of N6-methyladenosine and 5-methylcytosine RNA modifications and the respective enzymes that deposit (writer), remove (eraser) and facilitate (reader) function on viral infection. In this study we show how N4-acetyltransferase 10 (NAT10), a writer enzyme that acetylates cytidine to form N4-acetylcytidine (ac^4^C), affects Zika virus gene expression. Specifically, we found that NAT10 regulates viral infection kinetics by affecting the stability of select mRNAs involved in the innate immune response pathway. Our findings highlight another mode by which the innate immune response is robustly regulated in response to ZIKV infection.

## INTRODUCTION

Zika virus (ZIKV) is a re-emerged mosquito-transmitted flavivirus that in the last decade was associated with devasting outcomes (1–4). For example, during the outbreaks in the Americas the virus was associated with developmental and neurological issues babies following intrauterine infection (5–13). With the association of ZIKV with these anomalies paired with mosquito vector, mother-to-child and sexual transmission routes (14–16) there has been a rise in research efforts to investigate this understudied virus with the goal to develop new antiviral treatments or a vaccine.

The genome of ZIKV is a 10.8 kilobase (kb) single-stranded positive-sense RNA that closely resembles cellular mRNA (17). The ZIKV genome contains a 5’ methyl-guanosine cap, a single open reading frame (ORF), and 5’ and 3’ untranslated regions (UTR) but lacks a 3’ polyadenylated tail (18). The UTRs contain highly structured elements that function to promote translation, viral replication, and virion assembly (19). The UTRs also modulate cellular pathways such as the integrated stress and innate immune responses (20). The ORF encodes a single polyprotein that is proteolytically processed into three structural proteins (Capsid [C], precursor membrane [prM], envelope [E]) and seven non-structural proteins (NS1, NS2A, NS2B, NS3, NS4A, NS4B, and NS5) (18, 21). We and others recently showed that like cellular mRNAs, the nucleotides within the ZIKV genome may be modified with different chemical moieties (22, 23).

Chemical moieties or modifications may be deposited on RNA (24–30). These RNA modifications expand the breathtaking range of functions of RNA. For example, RNA modifications have been identified to affect RNA structure, stability and dynamics (31, 32), RNA splicing (33, 34), polyadenylation (35), transport (36), localization (37), and translation (29, 38–40). To date more than 170 RNA modifications are known (41). Indeed, RNA modifications can be added to all four nucleotides and onto different RNA molecules including viral RNA.

The N4-acetylcytidine (ac^4^C) modification is deposited on different RNA species by the ATP-dependent N-acetyltransferase 10 (NAT10) (42). The NAT10 writer enzyme has been shown to modify the 18S rRNA (43)which is impacts rRNA processing and ribosome biogenesis (44), and helps maintain the accuracy of mRNA translation (45). NAT10 together with the THUMP domain 1 (THUMPD1) co-factor acetylates cytidines in tRNAs (46). In *E. coli* tRNA^Met^, the first position of the anticodon loop is ac^4^C modified, while serine and leucine tRNAs contain ac^4^C modifications at other positions in the molecule (46). Recent next generation sequencing mapping of ac^4^C identified the modification within coding sequences of mRNAs which were shown to promote the translation efficiency and stability of mRNA (42).

Ac^4^C is also present on viral RNA. For example, ac^4^C marks on human Immunodeficiency virus type I (HIV-1) RNA, when silently mutated, decreased HIV-1 gene expression (47), and deletion of NAT10 the ac^4^C writer enzyme affected viral RNA stability (47). The viral RNA of Enterovirus 71 (EV71) was similarly shown to contain ac^4^C marks (48). Depletion (and deletion) of NAT10 and subsequent mutation of ac^4^C sites within the internal ribosome entry site (IRES) suppressed EV71 replication (48). Specifically, ac^4^C marks affected EV71 RNA stability (48). Moreover, EV71 ac^4^C modifications increased RNA translation via selective recruitment of the cellular poly-C binding protein 2 (PCBP2) to the IRES and increased the binding of RNA-dependent RNA polymerase to viral RNA (48). Ac^4^C marks have also been reported on the genomic RNA of Influenza A virus, although the biological significance of these marks was not investigated (49).

Here we investigated the role of the ac^4^C NAT10 writer enzyme on ZIKV gene expression. We found that in the absence of NAT10, ZIKV protein and RNA levels not only increased but the kinetics of the infection shifted earlier within the infection. We established that NAT10 impacted virus replication and turnover of the viral RNA, although there was no notable difference in the accumulation of the small flaviviral RNA. These data indicated that NAT10 likely regulated a cellular response. Indeed, we determined that NAT10 influenced the stability of select innate immune response mRNA transcripts. Overall, our data indicate that NAT10 plays a role in the activation of the host innate immune response by acting as the acetylator for immune factors and upstream activating factors.

## RESULTS

### NAT10 restricts the abundance of ZIKV protein and RNA but has no effect on the production of infectious virions at 48 hours post-infection

We previously used a highly sensitive and unbiased approach to broadly identify RNA modifications on different positive-sense RNA viruses (22). Our mass spectrometry analysis showed that the global RNA modification profile changed during viral infection. We found no difference in the abundance of single modifications on all cytidines between mock- and ZIKV-infected cells (22). Interestingly however, in ZIKV-infected cells the overall levels of N4-acetylcytidine (ac^4^C) decreased (22). In that study we also examined the distribution of RNA modifications on ZIKV RNA using RNA affinity pulldowns of intracellular and extracellular viral RNA prior to mass spectrometry analysis. We identified 34 different RNA modifications on ZIKV RNA (22). In addition to N6-methyladenosine (m^6^A), which was previously reported on ZIKV RNA (50, 51), we identified pseudouridine as the most abundant chemical moiety on ZIKV RNA followed by ac^4^C (22). To begin to understand the significance of ac^4^C, in this study we investigated the role of NAT10, the ac^4^C writer enzyme, on ZIKV infection. Specifically, HeLa wild-type (WT) and NAT10 knockout (KO) cells were either mock-infected or infected with ZIKV, and viral and cellular protein and RNA, and virions released into the cell culture media harvested at 48 hours post-infection. By western blot we observed that the levels of NAT10 were only faintly detectable in the NAT10 KO cells (Figure 1A). Of note, although the HeLa NAT10 KO cell line was created using the CRISPR-Cas9 gene deletion strategy, through low-levels of alternative splicing some NAT10 expression remains (52). An examination of THUMPD1, the cofactor that assists with acetylating tRNA (46), showed a decrease in abundance in the HeLa NAT10 KO cells (Figure 1A). When we examined the levels of ZIKV NS1 protein we observed a two-fold increase in the amount of viral protein (Figure 1A, 1B). RT-qPCR analysis of the levels of ZIKV RNA similarly showed a significant increase in NAT10 KO cells as compared with the WT cells (Figure 1C). These data suggest that NAT10 (and possibly ac^4^C marks on the viral RNA) affects the abundance of ZIKV protein and RNA in cells. Despite the increase in ZIKV protein and RNA, when we performed plaque assays to quantify the amount of infectious virus released into the media, we found no difference in ZIKV titers from infected WT and NAT10 KO cells (Figure 1D). Together these data indicate that NAT10 restricts the accumulation of ZIKV protein and RNA levels, without impacting the abundance of infectious virions.

**Figure 1:**
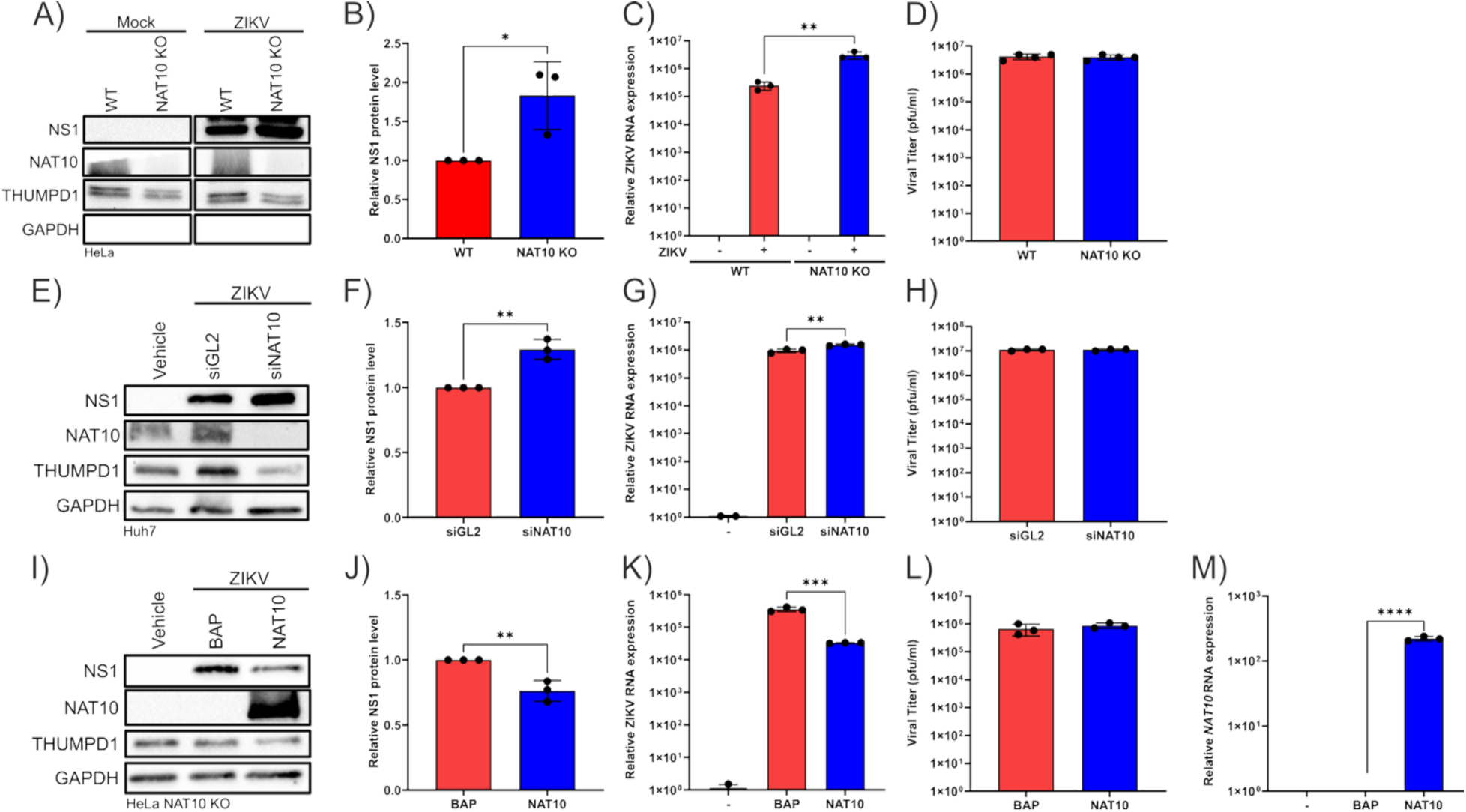
NAT10 differentially affects the levels of ZIKV protein and RNA but not viral titers. (A-D) HeLa wildtype (WT) and NAT10 knockout (KO) cells were either mock-infected or infected with ZIKV at MOI of 5 pfu/cell, harvested at 48 hours post-infection and analysis of viral and cellular proteins and RNA, and ZIKV titers was undertaken. (A, E, I) Western blots show the abundances of ZIKV NS1 protein, and cellular proteins NAT10, THUMPD1 and GAPDH in mock- and ZIKV-infected cells. (B, F, J) ZIKV protein abundance was semi-quantified by determining the band density of ZIKV NS1 and GAPDH using Image Lab. The levels of NS1 protein are represented relative to GAPDH. (C, G, K) RT-qPCR showing levels of ZIKV RNA relative to *ACTB* mRNA. (D, H, L) ZIKV titers (pfu/ml) following infection in WT or NAT10 KO cells was determined by plaque assay. (E-H) Huh7 cells were transfected with Lipofectamine 3000 (vehicle), a nonspecific control siRNA targeting *Gaussia* luciferase (siGL2) or a siRNA targeting *NAT10* mRNA. Forty-eight hours following transfection of the siRNAs, cells were infected with ZIKV (MOI = 5 pfu/ml) and viral gene expression examined 48 hours post-infection. (E) Western blot validates NAT10 knockdown and the effect on ZIKV NS1 protein levels. The abundance of THUMPD1 and GAPDH are also shown with GAPDH as a loading control. (F) Semi-quantifications of ZIKV NS1 levels relative to GAPDH. (G) Analysis of ZIKV RNA levels relative to *ACTB* mRNA as determined by RT-qPCR. (H) Plaque assays report the amount of infectious virus produced in Huh7 control and NAT10 siRNA depleted cells. (I-M) The effect of rescuing NAT10 protein on ZIKV was investigated by transfecting HeLa NAT10 KO cells with Lipofectamine 3000 reagent (vehicle), p3xFlag-BAP and p3xFlag-NAT10 plasmids. Twenty-four hours post-transfection, cells were infected with ZIKV (MOI = 5 pfu/ml) followed by analysis of viral and cellular proteins and RNA, and ZIKV titers. (I) Western blots show the abundance of ZIKV NS1 protein, and cellular proteins NAT10, THUMPD1 and GAPDH in mock- and ZIKV-infected cells. (J) Semi-quantification of the ZIKV NS1 protein relative to GAPDH. (K) RT-qPCR showing the amount of ZIKV RNA relative to *ACTB* mRNA. (L) ZIKV titers (pfu/ml) following infection in WT or NAT10 KO cells transfected with control and NAT10 plasmids. (M) RT-qPCR showing levels of NAT10 RNA relative to *ACTB* mRNA. The data are representative of at least three independent experiments. Error bars show ± SD. Statistical analysis was performed using an unpaired Student t-test; *P< 0.05, **P < 0.01. Unannotated data were found to be not significant. Quantitative data from HeLa WT and NAT10 KO cells are reported in red and blue, respectively

To validate these findings, we next performed an orthogonal assay using a control siRNA targeting *Guassia* luciferase (siGL2) and a NAT10 target specific siRNA. The siRNAs were transfected into Huh7 cells, infected with ZIKV and viral gene expression examined. By western blot we observed that the levels of NAT10 were depleted in the Huh7 cells transfected with siNAT10 (Figure 1E). Consistent with the expression in the HeLa NAT10 KO cells, the expression of THUMPD1 was also reduced when NAT10 was knocked down (Figure 1E). Likewise, the levels of ZIKV NS1 protein increased two-fold in Huh7 cells transfected with the NAT10 specific siRNA compared to the control siRNA (Figure 1 E, 1F). An examination of the abundance of ZIKV RNA by RT-qPCR similarly showed a significant increase in siNAT10 transfected cells as compared siGL2 transfected Huh7 cells (Figure 1G). By plaque assay we again observed no difference in ZIKV titers from NAT10 depleted cells compared to control (Figure 1H). These data support the results from the HeLa Nat10 KO cells and demonstrate that the role of NAT10 during ZIKV infection is not cell type specific.

To further explore the role of NAT10, we investigated the effect of rescuing NAT10 in HeLa NAT10 KO cells. Given our previous results with NAT10 KO and siNAT10, we predicted that ZIKV protein and RNA levels would decrease when NAT10 expression was restored. To this end, we first transfected HeLa NAT10 KO cells with either a control plasmid expressing 3xFlag-tagged Bacterial Alkaline Phosphatase (3xFlag-BAP) or 3xFlag-tagged NAT10 and then infected with ZIKV. By western blot, we observed that NAT10 protein was absent in NAT10 KO cells transfected with either transfection reagent alone (Vehicle) or 3xFlag-BAP and overexpressed in 3xFlag-NAT10 transfected cells (Figure 1I). *NAT10* mRNA levels significantly increased in NAT10 KO cells that were transfected with 3xFlag-NAT10 (Figure 1M). The levels of the THUMPD1 cofactor protein modestly decreased, albeit not significantly (data not shown), when NAT10 was overexpressed (Figure 1I). As predicted, the levels of ZIKV NS1 protein significantly decreased when NAT10 expression was restored in the NAT10 KO cells, compared to cells transfected with either transfection reagent alone or the control plasmid (Figure 1I,J). RT-qPCR analysis similarly showed a significant decrease in the amount of ZIKV viral RNA in NAT10 KO cells expressing 3xFlag-tagged NAT10 as compared with the cells expressing the control plasmid (Figure 1K). Similar to ZIKV infection in the HeLa NAT10 KO cells, rescue of NAT10 did not significantly impact the production of ZIKV virions at 48 hours post infection (Figure 1L). Together, these data show that the levels of ZIKV protein and RNA are impacted by NAT10 abundance.

### THUMPD1, the NAT10 co-factor, does not affect ZIKV gene expression

In NAT10 HeLa and NAT10 knockdown Huh7 cells, we observed that the levels of THUMP D1 were decreased (Figure 1A). Given the effect of NAT10 on ZIKV protein and RNA levels, we wondered if the decrease in THUMPD1 also contributed to the differential effects on ZIKV. To examine this, we used CRISPR-Cas9 gene editing to knockout THUMPD1 in HeLa cells. Next, HeLa WT and THUMPD1 KO cell lines were either mock infected or infected with ZIKV and viral protein, RNA and titers examined. By western blot, we observed a clear knockout of the THUMPD1 protein (Figure 2A). RT-qPCR also demonstrated a significant decrease in *THUMPD1* mRNA levels in both mock- and ZIKV-infected HeLa THUMPD KO cells compared to WT cells (Figure 2B). The levels of NAT10 protein were similar in both WT and THUMPD1 KO cells, and in the absence or presence of ZIKV infection (Figure 2A). Although knockout of NAT10 affects ZIKV protein and RNA levels, deletion of THUMPD1 did not affect ZIKV gene expression (Figure 2). Thus, the protein that binds tRNA to facilitate the acetylation of serine and leucine tRNAs by NAT10 does not have a role in ZIKV infection.

**Figure 2:**
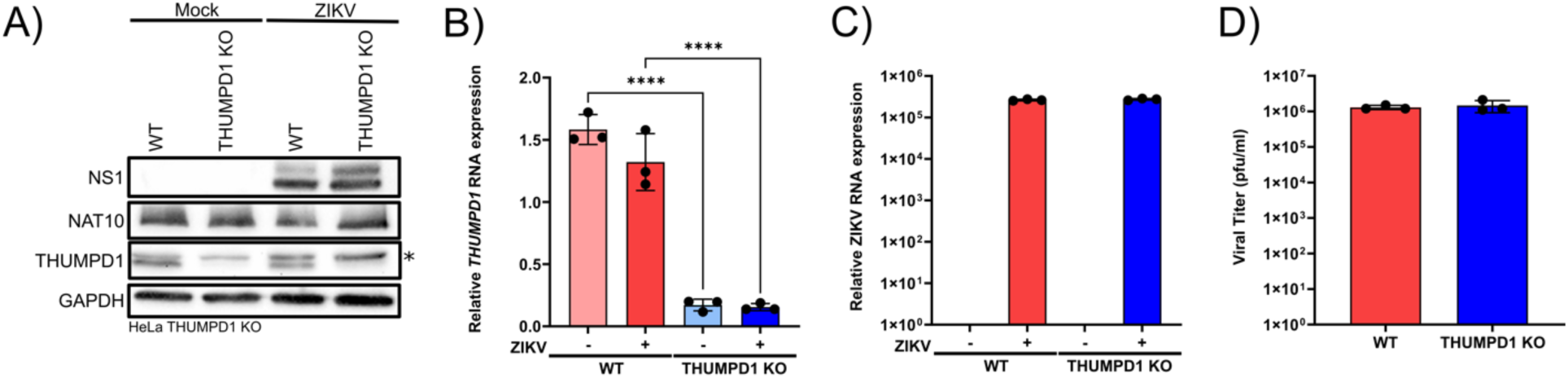
ZIKV gene expression is not significantly affected by THUMPD1, a cofactor of NAT10. CRISPR-Cas9 gene editing was used to knock out the expression of THUMPD1 in HeLa cells. HeLa WT and THUMPD1 KO cells were mock- or ZIKV-infected at a MOI of 5 pfu/ml and protein, RNA and viral titers examined 48 hours post-infection. A) Western blots showing the levels of ZIKV NS1 protein, and cellular NAT10, THUMPD1 and GAPDH protein levels. GAPDH was included as a loading control for the Western blot. *Denotes a nonspecific band. B) Semi-quantification of the western blot in panel A determining the abundance of ZIKV NS1 protein relative to GAPDH. RT-qPCR analysis of B) *THUMPD1* mRNA transcript and C) ZIKV RNA. The data report the abundance of each cellular and viral RNA relative to *ACTB*. D) Quantification of ZIKV infectious virions in the media as determined by plaque assay. All data are representative of at least three independent experiments. Error bars represent ± SD. An unpaired Student t-test was used to determine statistical significance. ****P< 0.0001. Unannotated data were found to be not significant. Quantitative data from HeLa WT and THUMPD1 KO cells are shown in red and blue, respectively.

### ZIKV infection kinetics are impacted by NAT10

Given that ZIKV protein and RNA abundances increase in NAT10 knockout and depleted cells (Figure 1) we posited that this increase in ZIKV gene expression was likely due to a change in infection kinetics or the result of increased production of viral protein and RNA. To elucidate this, we first compared ZIKV protein, RNA and titers over time from HeLa WT and NAT10 KO cells. By western blot, the levels of ZIKV NS1 protein in HeLa WT cells were first detectable at 36 hours post-infection (Figure 3A). In contrast, in the HeLa NAT10 KO cells ZIKV NS1 was detectable at 24 hours post-infection (Figure 3B). An examination of the abundance of ZIKV RNA by RT-qPCR similarly showed an increase in NAT10 KO cells beginning at the 18-hour timepoint as compared with the WT cells (Figure 3E). These data show that NAT10 (and possibly ac^4^C marks on the viral RNA) affect ZIKV protein and RNA abundance in cells by impacting the infection kinetics.

**Figure 3:**
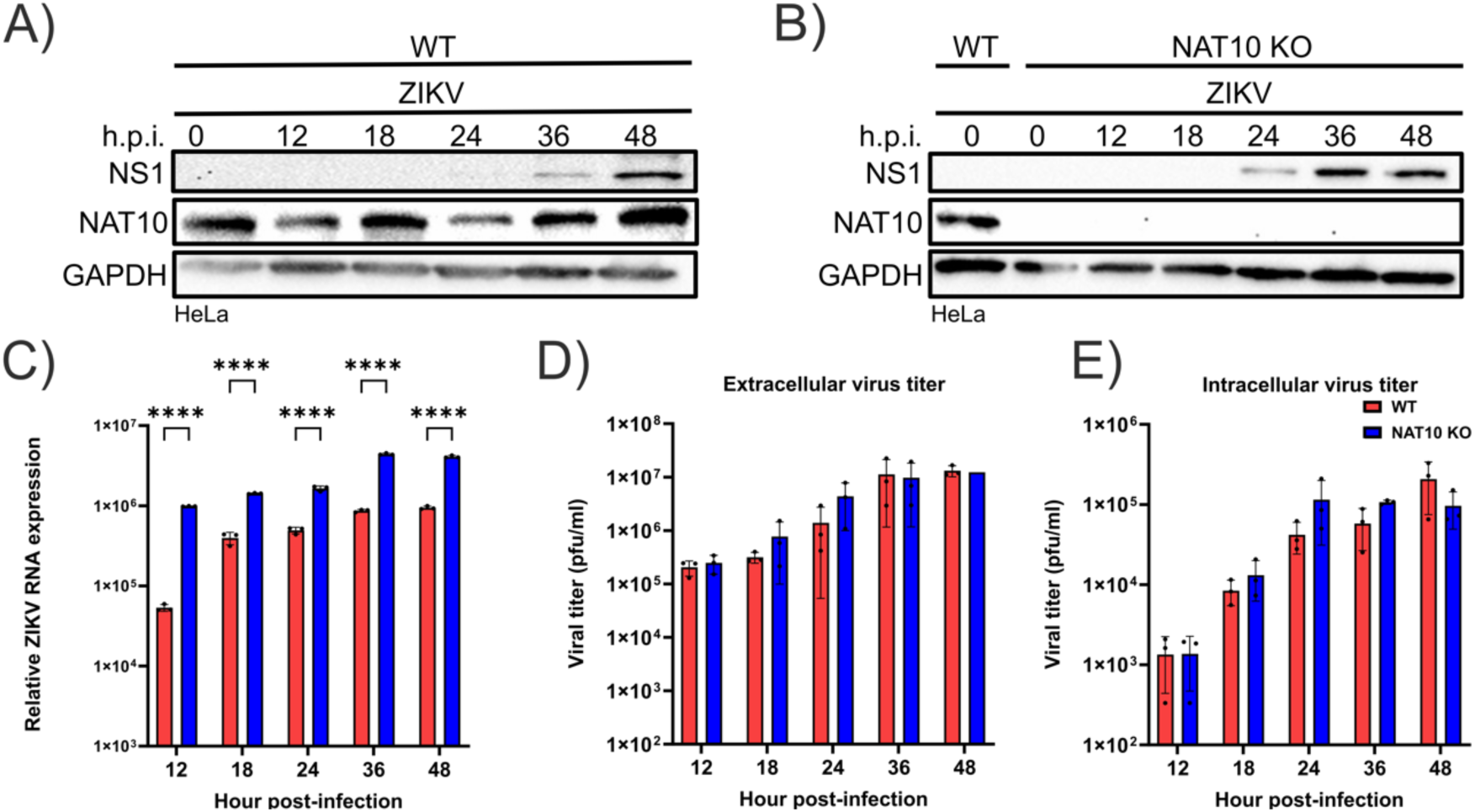
NAT10 influences the kinetics of ZIKV infection. HeLa WT and NAT10 KO cells were mock-infected or infected with ZIKV at MOI of 5 pfu/cell. At 12-, 18-, 24-, 36-, and 48-hours post-infection (h.p.i.) cells were harvested for analysis of viral and cellular proteins and RNA levels, and quantification of intracellular and extracellular ZIKV titers. Western blots in A) HeLa WT and B) HeLa NAT10 KO cells at each respective time point show the abundances of ZIKV NS1, and cellular NAT10 and GAPDH proteins. C) RT-qPCR was performed to determine the amount of intracellular ZIKV RNA relative to *ACTB* mRNA. Plaque assays were performed to quantify the amount of ZIKV produced from HeLa WT and NAT10 KO infected cells. Viral titers in D) the media (extracellular titer) and E) infected cells (intracellular titer) are reported. The experiments were undertaken three independent times. Error bars show ± SD. Statistical analysis was performed using 2-way ANOVA. ****P<0.0001. Unannotated data were found to be not significant. Quantitative data from HeLa WT and NAT10 KO cells are presented in red and blue, respectively.

Given the earlier detection of ZIKV protein in the HeLa NAT10 cells, we speculated that we might also observe a difference in viral titers at an earlier timepoint. An examination of the virions released into the media showed comparable levels between WT and NAT10 KO cells at 12 hours post-infection (Figure 3D). At 19- and 24 hours post-infection, the viral titers in the NAT10 KO cells were higher, albeit not significantly (Figure 3D). At 36- and 48 hours post-infection, we again found no significant difference in ZIKV extracellular titers from infected WT and NAT10 KO cells (Figure 3C). We also assayed intracellular RNA titers, with the reasoning that NAT10 might influence virion trafficking thus restricting the release of assembled virions from the cell. Like the plaque assays with the extracellular virions, intracellular titers were modestly higher at 18- to 36- hours post-infection (Figure 3D). Overall, these data show that ZIKV protein and RNA accumulate earlier in the NAT10 KO cells, but the difference in the kinetics of ZIKV gene expression does not impact virus assembly and release.

### NAT10 affects the translation, replication and turnover of ZIKV RNA

Given that NAT10 affected the kinetics of ZIKV protein and RNA, we next examined the role of NAT10 on ZIKV translation and/or replication. To investigate this, we used a subgenomic *Renilla* luciferase replicon that contains a T7 promoter, 5’ and 3’ UTRs flanking an open reading frame that encodes the first 22 amino acids of capsid, *Renilla* luciferase and the nonstructural proteins NS1-NS5 (53). Following *in vitro* transcription, capping and transfection of the viral replicon RNA, we measured luciferase activity as a proxy for translation and replication. Specifically, we assayed translation at 4- and 8-hours post-transfection into either HeLa WT and NAT10 KO cells, and replication at 24-, 36- and 48-hours post-transfection (Figure 4A). From this assay we determined that initial translation was higher in the HeLa NAT10 KO cells compared to WT cells at 4-hours post-transfection, but not at 8-hours, albeit not statistically significant (Figure 4A). Notably, however, in the absence of NAT10 replication at 24-, 36- and 48-hours was significantly higher (Figure 4A). Indeed, when we assayed luciferase activity between 8- and 20-hours post-transfection, replication of the replicon RNA increased earlier and at higher levels in the NAT10 KO cells compared to WT (Figure 4B). These data are consistent with the increased protein and RNA levels determined during ZIKV infection (Figure 3A-3C). As an additional control for replication, we assayed a replication-deficient replicon in which the GDD motif in the NS5 active site was mutated to GNN (53), rendering the ZIKV NS5 RNA-dependent RNA polymerase inactive (Figure 4B). Consistent with the replication competent construct, translation of the replication incompetent replicon was higher at 4-hours post-transfection in the NAT10 KO cells compared to WT cells (Figure 4C). Unexpectedly however the luciferase signal appeared to decrease more in the NAT10 KO cells than in WT cells (Figure 4C), suggesting the replicon RNA might be turned over sooner in the absence of NAT10. To investigate the effect of NAT10 on the decay of the ZIKV replicon RNA, we assayed luciferase activity from the replication incompetent replicon between 8- and 20-hours post-transfection (Figure 4D). In contrast to the expression of the replication competent replicon RNA (Figure 4A and Figure 4B), translation of the replication incompetent ZIKV replicon was higher in the NAT10 KO cells compared to WT (Figure 4 and Figure 4D). In HeLa WT cells, the luciferase activity from the replication incompetent RNA gradually decreased from 8-hours post-transfection (half-life of ZIKV RNA = 18.1 hours) (Figure 4D). In the HeLa NAT10 KO cells, the replicon RNA precipitously decreased between 8- and 14-hours post-transfection (Figure 4D). The half-life of the replicon RNA in the HeLa KO cells was

**Figure 4:**
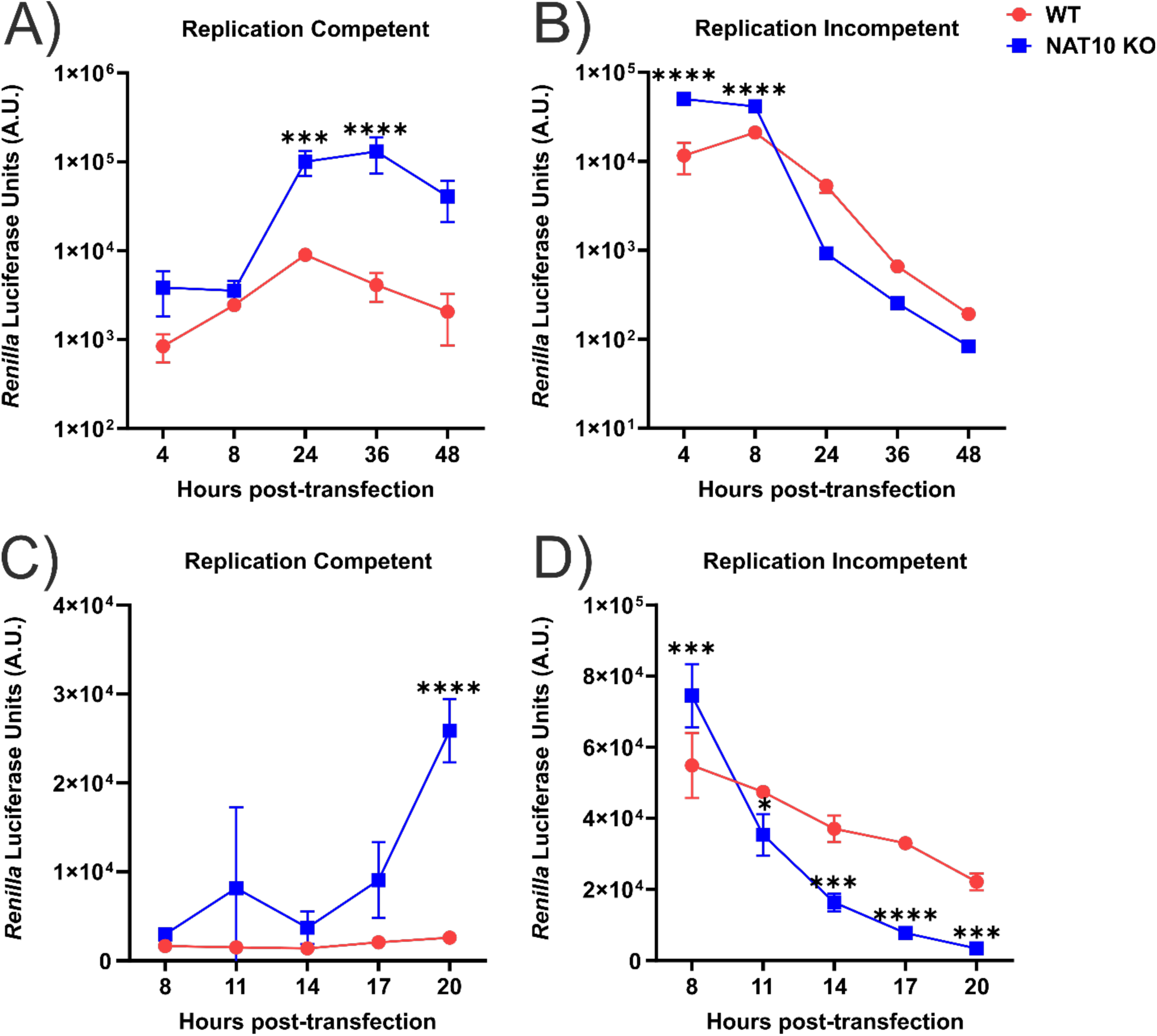
NAT10 affects ZIKV replication and decay of the viral RNA. *In vitro* transcribed and capped subgenomic ZIKV reporter (*Renilla* luciferase, RLuc) replicon RNA was prepared, transfected into HeLa WT (red) and NAT10 KO (blue) cells and harvested at the time points reported on the respective x-axes. Panel A and C show data from the replication competent ZIKV replicon. Panels B and D report data from cells transfected with the replication-incompetent ZIKV replicon. The replication-incompetent ZIKV replicon harbors a GDD to GNN mutation in the active site of NS5 RNA-dependent RNA polymerase rendering the enzyme replication inactive. A, B) Analysis of the translation and replication of A) replication-competent and B) replication-incompetent ZIKV replicons 4-48 hours post-transfection. *Renilla* luciferase units were measured in arbitrary units (A.U.) and are a proxy for ZIKV translation and replication. Four and 8-hours post-transfection report on translation of the ZIKV replicon RNA, and replication of ZIKV RNA was assayed at 24-, 36- and 48-hours post-transfection. Using the decrease of the *Renilla* luciferase activity of the replication incompetent replicon as a readout for ZIKV RNA decay, the half-life (t_1/2_) of the RLuc replicon in HeLa WT and NAT10 KO cells was calculated as 15.7 ± 1 hours and 10.8 ± 0.2 hours, respectively. C, D) *Renilla* luciferase activity from C) replication-competent and D) replication-incompetent ZIKV replicon RNAs measured at 8-, 11-, 14-, 17- and 20-hours post-transfection. Using the decrease of the *Renilla* luciferase activity of the replication incompetent replicon as a readout for the decay ofZIKV RNA, the half-life (t_1/2_) of the RLuc replicon in HeLa WT and NAT10 KO cells was calculated as18.1 ±1 hours and 10.8 ±.34 hours, respectively. The data are representative of three independent experiments and errors bars represent ± SD. Statistical significance of the assay data from HeLa WT and NAT10 KO cells at each time point was determined using an unpaired Student *t*-test. *P<0.05, ***P<0.001, ****P<0.0001, and not significant data are not annotated.

### 10.7 hours

The examination of the expression ZIKV replicon RNA suggests that NAT10 affects both turnover and replication of ZIKV RNA. Given these seemingly opposing effects, we posited that in the absence of NAT10, the rapid turnover of the replicon RNA would alter the levels of small flaviviral (sf) RNAs (either appearing earlier or at higher levels) compared to WT cells. The cellular 5’-to-3’ exonuclease XRN1has been shown to incompletely degrade flaviviral RNAs to produce sfRNAs, or viral 3’ UTRs that are resistant to XRN1 decay, which in turn have been reported to modulate the immune response to permit higher levels of ZIKV replication (54). Given that the immune response in HeLa cells is not robust, prior to testing our hypothesis, we used CRISPR-Cas9 gene editing to knockout NAT10 in the A549 human lung carcinoma cell line. Although ZIKV in humans does not cause lung disease, we and others have previously shown that A549 cells have a robust immune response to *flavivirus* infection (55–58). We have also shown that changes in differential gene expression and alternative splicing transcripts in ZIKV-infected A549 cells mirror those in PBMCs isolated from ZIKV infected patients (59).

We first validated the expression of the replication competent and incompetent ZIKV luciferase replicon in A549 WT and NAT10 KO cells (Figure 5A, 5B). Consistent with our earlier experiment in HeLa cells we found that replication is initiated earlier and at higher levels in the A549 NAT10 KO cells compared to WT (Figure 5A). Likewise, the replicon RNA decays more rapidly between 8- and 11 hours post-transfection in A549 cells deficient of NAT10 compared to WT (Figure 5B). Of note, by 20-hours post-transfection, *Renilla* luciferase activity from remaining replication incompetent replicon RNA was comparable (Figure 5B). To examine the production of sfRNA, we infected A549 WT and NAT10 KO cells with ZIKV, and isolated total RNA at 12-, 18-, and 24-hours post-infection. Total RNA was separated in a denaturing formaldehyde agarose gel, and 28S rRNA and ZIKV sfRNA visualized by northern blotting. Quantification of the relative amount of ZIKV sfRNA compared to 28S rRNA, over three independent experiments, showed no statistical difference between the A549 WT and NAT10 KO cells (Figure 5C). Thus, despite the rapid turnover of the ZIKV *Renilla* luciferase replicon in NAT10 KO cells (Figure 4D and Figure 5B), the accumulation of sfRNA was not affected (Figure 5C). These data suggest that a putative regulation of ZIKV gene expression by NAT10 is likely not mediated by the viral sfRNA and viral directed sequestration of immune factors.

**Figure 5:**
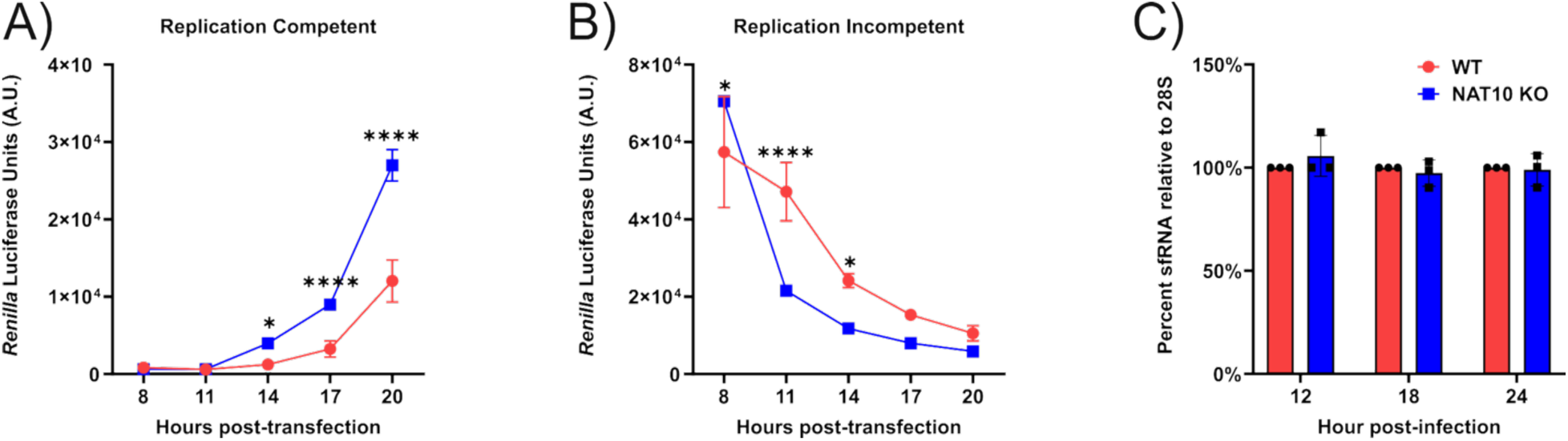
NAT10 influences the stability of ZIKV RNA but not the production of ZIKV small flaviviral (sf) RNAs. A, B) *In vitro* transcribed and capped subgenomic ZIKV RLuc replicon RNA was prepared and transfected into A549 WT and NAT10 KO cells. *Renilla* luciferase activity, in arbitrary units (A.U.), from the A) replication competent and B) replication incompetent ZIKV reporter replicon was measured at 8-, 11-, 14-, 17- and 20-hours post-transfection. ZIKV RNA half-life in A549 WT and NAT10 KO cells was calculated as 13.4 ±1 hours and 10.1 ± 0.4 hours, respectively. C) A549 WT and NAT10 KO cells were infected with ZIKV (MOI = 5 pfu/cell) and harvested at 12-,18-, and 24 hours post-infection. Total RNA was extracted and sized in a 1% denaturing formaldehyde agarose gel. From the northern blot the abundance of ZIKV sfRNA was quantified relative to the 28S rRNA using Image Lab software. The abundance of sfRNA in A549 WT cells was set to 100% and the sfRNA levels in NAT10 KO cells are shown relative to WT. The data are representative of three independent experiments. Error bars show ± SD. Statistical significance was interrogated using a 2-way ANOVA. *P<0.05 and ****P<0.0001. Unannotated data are not significant. The abundance of the sfRNA at each time point was determined to be not significant.

### NAT10 affects the expression of components associated with the innate immune response

We next examined if the increase in ZIKV replication (Figures 4A, 4B and Figure 5A), even following the apparent turnover of viral RNA (Figure 4D and Figure 5B), might be a result of an attenuation of the innate immune response. To examine this, we either mock- and ZIKV-infected A549 WT and NAT10 KO cells and examined protein and RNA abundances and viral titers at 24- and 48-hours post-infection (Figure 6). Consistent with ZIKV infection in the HeLa NAT10 KO cells, ZIKV NS1 increased at 24 hours post-infection in the A549 NAT10 KO cells compared to WT cells and was comparable at 48 hours post-infection (Figure 6A). Notably, in the absence of NAT10, ZIKV RNA levels were higher at both 24- and 48 hours post-infection (Figure 6B) yet viral titers were only increased at 24 hours post-infection (Figure 6C). These data support our earlier conclusion that NAT10 regulates ZIKV gene expression.

**Figure 6:**
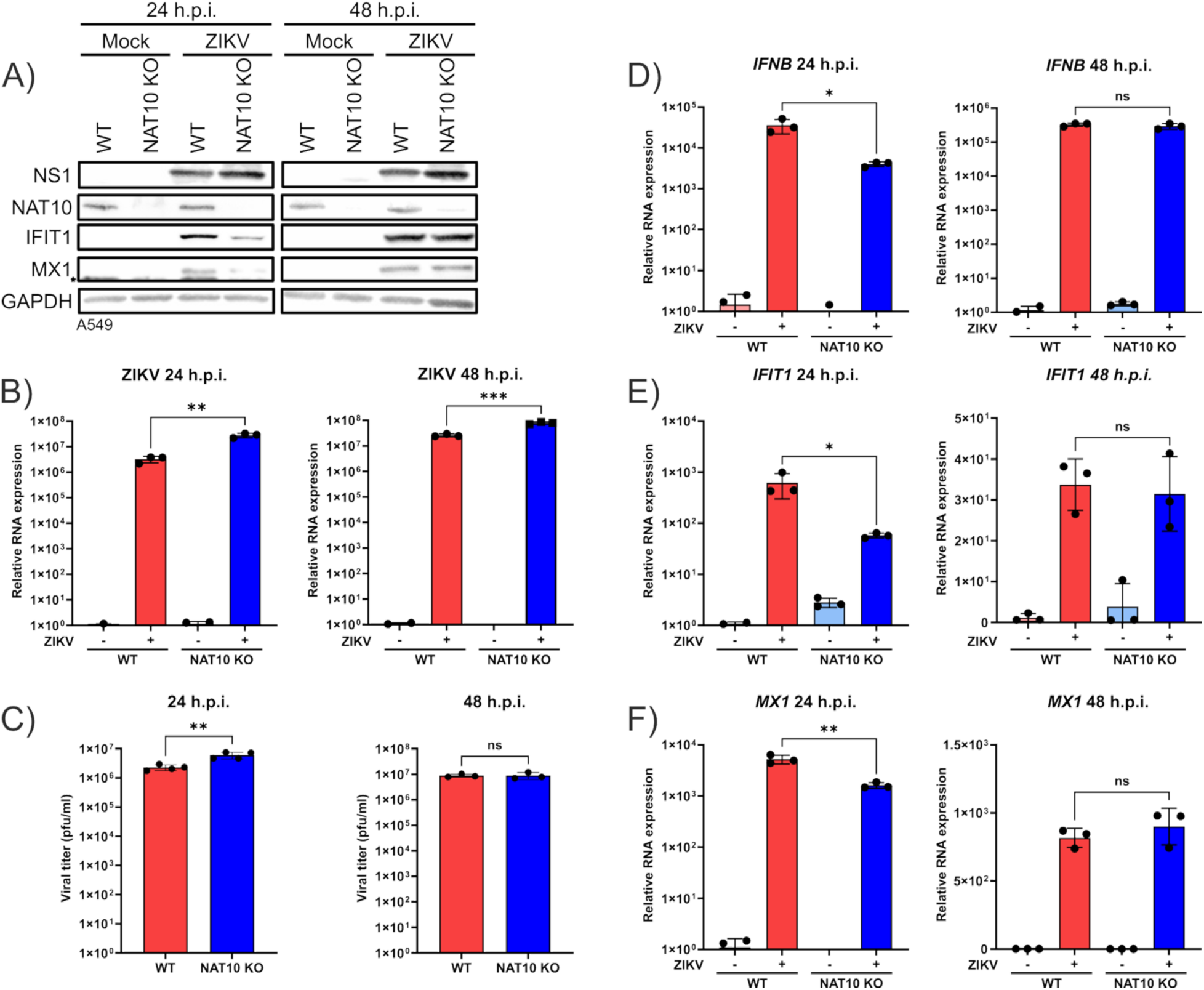
NAT10 affects the abundance of select components associated with the innate immune response pathway at 24 hours post-infection. A549 WT and NAT10 KO cells were either mock-infected or infected with ZIKV (MOI = 5 pfu/cell) and cells were harvested at 24- and 48-hours post-infection. A) The levels of ZIKV NS1, and cellular proteins NAT10, MX1, IFIT1 and GAPDH at 24- and 48-hours post-infection (h.p.i.) were examined by western blot analysis. GAPDH was used as a loading control for the assay. * MX1 (and IFIT1) proteins at 24 hours post-infection were probed after detection of NAT10. The nonspecific band denoted by an astericks is from the detection of NAT10. B) RT-qPCR was used to quantify ZIKV RNA abundance relative to *ACTB* at each time point in both mock- and ZIKV-infected WT and NAT10 KO cells. C) Plaque assays were undertaken to determine ZIKV titer in infected WT and NAT10 KO A549 cells. D-F) RT-qPCR was performed to determine the levels of *IFNB*, *IFIT1,* and *MX1* transcripts at 24- and 48 hours respectively in A549 WT and NAT10 KO mock- and ZIKV-infected cells. The levels of each transcript are reported relative to *ACTB*. The data are representative of three independent experiments. Error bars represent ± SD. Statistical analysis was performed using an unpaired Student *t*-test. *P<0.05, **P<0.01, ***P<0.001, and ns – not significant.

Next, we investigated ZIKV induction of the innate immune response in A549 WT and NAT10 KO cells. By western blotting at 24 hours post-infection, we observed that abundance of IFIT1 and MX1, two well established interferon (IFN) stimulated proteins, was decreased in the NAT10 KO cells (Figure 6A). Consistent with our other data, however, by 48 hours post-infection the IFIT1 and MX1 protein abundances were comparable at 48 hours post-infection in both WT and NAT10 KO cells (Figure 6A). RT-qPCR analysis of interferon-β (*IFNB*), *IFIT1* and *MX1*, similarly showed decreased transcript levels at 24 hours post-infection in NAT10 KO cells compared to WT cells (Figure 6D-6F). Indeed, the reduced expression of both innate immune response mRNAs and proteins likely contributes to the increased ZIKV gene expression levels at 24 hours post-infection in cells deficient of NAT10 (Figure 6A-6C). Notably, in both WT and NAT10 KO cells ZIKV-infected cells, we observed no statistically significant difference in the innate immune response transcripts induced in ZIKV infected cells at 48 hours post-infection (Figure 6D-6F). The comparable expression of these genes at 48 hours post-infection might in part influence the equivalent levels viral protein and titers at the 48-hour time point (Figure 6A-6C). Together these data suggest that early during infection the increased ZIKV protein and RNA levels are likely a consequence of a dampening activation and expression of genes within the innate immune response pathway.

### The effect of NAT10 on ZIKV gene expression occurs via activation of the JAK/STAT signaling pathway and modulation of ISGs

*IFNB* mRNA and IFIT1 and MX1 protein and RNA levels are reduced in ZIKV-infected NAT10 KO cells (Figure 6). From these data, the decreased expression of IFIT1 and MX1 might be a consequence of reduced transcription, mRNA turnover or dampened JAK/STAT signaling resulting from decreased *IFNB* expression. To investigate the effect on the JAK/STAT signaling pathway, A549 WT and NAT10 KO cells were either mock-infected or infected with ZIKV and then treated with Ruxolitinib, a pharmacological compound that selectively inhibits JAK1 and JAK2, two key tyrosine kinases in the JAK/STAT signaling pathway (60, 61). We hypothesized that the negative effect of NAT10 deletion on reduced *IFNB* levels, and the positive effect on ZIKV gene expression, would be negated by Ruxolitinib and blocking the JAK/STAT signaling cascade. Thus, ZIKV gene expression in Ruxolitinib-treated WT and NAT10 KO cells would be comparable. By western blot we observed that the abundance of STAT1 was decreased in NAT10 KO cells compared to WT cells, although STAT1 activation via phosphorylation (p-STAT1) was similar in both ZIKV-infected and DMSO-treated WT and KO cells (Figure 7A). Likewise, and consistent with our earlier data (Figure 6), the abundances of IFIT1 and MX1 proteins were also reduced in NAT10 KO cells (Figure 7A). Treatment with the JAK inhibitor did not affect overall STAT1 protein levels, however the resulting inhibition of JAK and downstream signaling resulted in p-STAT1, IFIT1 and MX1 proteins being not detectable by western blot (Figure 7A). *STAT1*, *IFIT1* and *MX1* mRNA levels mirrored the observed effects on protein abundances in A549 WT and NAT10 KO in mock- and ZIKV-infected cells without and with the JAK inhibitor (Figure 7D-7F). In NAT10 KO cells treated with DMSO, ZIKV protein and RNA levels, and titers were increased compared to WT cells (Figure 7A-7C). However, when WT and NAT10 KO cells were infected and treated with JAK1/2 inhibitor, ZIKV gene expression showed no statistical difference (Figure 7A-7C). Overall, these data suggest that NAT10 has a role in regulating at least some components of the innate immune response pathway.

**Figure 7:**
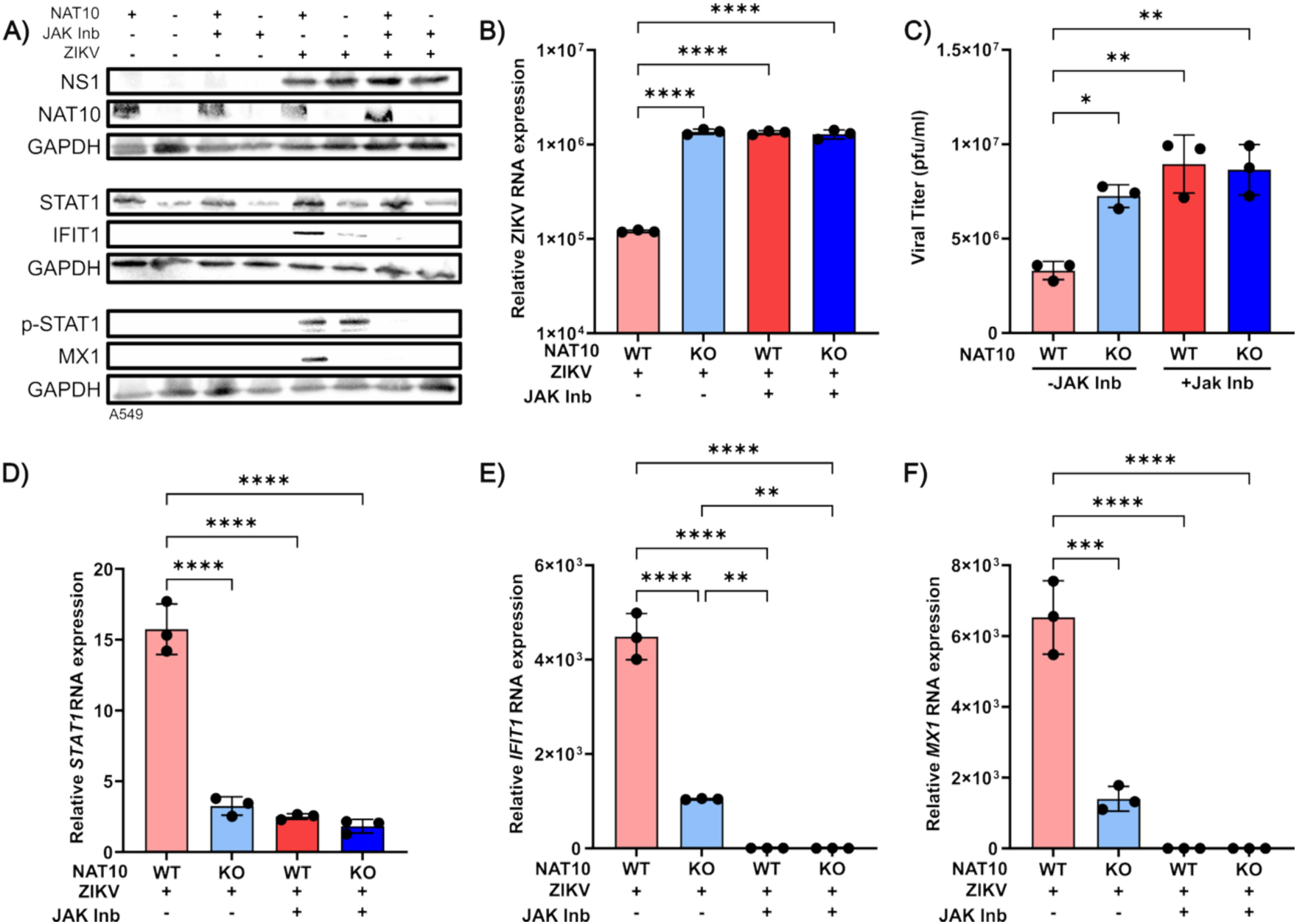
Inhibition of the JAK/STAT signaling pathway limits the effect of NAT10. A549 WT and NAT10 KO cells were preincubated with PBS or ZIKV (MOI = 5 pfu/cell) for 1 hour at 37°C. Hereafter, media containing either DMSO or 30 nM of Ruxolitinib, a JAK1/2 inhibitor (JAK Inb), was replaced on mock- and ZIKV-infected cells. Viral and cellular proteins and RNA, and viral titers were assayed 24 hours post-infection. A) Western blots showing the abundance of ZIKV NS1 and NAT10 proteins. Components of the innate immune response namely STAT1, phosphorylated STAT1 (p-STAT1) and interferon stimulated proteins IFIT1 and MX1 are also shown. B-E) show data from ZIKV-infected A549 WT and NAT10 KO cells. Analysis of Mock-infected cells is not included. B) Quantification of ZIKV RNA levels by RT-qPCR relative to *ACTB* mRNA. C) ZIKV titers were determined by plaque assay. RT-qPCR analysis of the fold change of D) *STAT1*, E) *IFIT1* and F) *MX1* mRNAs. Quantitative data from ZIKV infection in A549 WT and NAT10 KO cells are shown as red and blue respectively. Contrasting color shade indicates without and with JAK1/2 inhibitor (Ruxolitinib). Data in this figure are representative of three independent experiments, and errors bars indicate ± SD. A 2-way ANOVA was used to determine statistical significance. **P<0.01, ***P<0.001 and ****P<0.0001.

### Interferon stimulated IFIT1 and MX1 mRNAs decay more rapidly in the absence of NAT10

In NAT10 KO cells, ZIKV gene expression is increased in part due to the reduced expression of *IFNB* and other components of the innate immune response pathway (Figure 6 and Figure 7). Notably, STAT1 protein levels are decreased in NAT10 KO cells yet p-STAT1 levels were comparable (Figure 7A). Dang and colleagues previously reported that during Sindbis virus infection, NAT10 altered the mRNA stability of lymphocyte antigen six family member E (LY6E), an innate immune response factor (62). To investigate mRNA turnover, A549 WT and A549 NAT10 KO cells were first infected with ZIKV and then at 16 hours post-infection cells were treated with Actinomycin D, and harvested at 0-, 1-, 2-, 3-, 4-, 6-, and 8-hours post-treatment. Total cellular RNA was isolated and the abundance of select mRNA transcripts were assayed by RT-qPCR (Figure 8). In examining the amount of *STAT1*, *IFIT1* and *MX1* mRNAs remaining, we observed increased turnover of the transcripts (Figure 8A-8C). For example, the half-life of *STAT1* mRNA in NAT10 KO cells decreased from 18.5 hours to 10 hours compared to WT cells (Figure 8A). Indeed, the decrease in *STAT1* mRNA stability was also reflected in the reduced amount of STAT1 protein in both mock- and ZIKV-infected cells (Figure 7A). Additionally, the decreased *STAT1* mRNA stability did not change when the JAK/STAT signaling cascade was inhibited with Ruxolitinib, showing *STAT1* turnover was not regulated by activation (or inhibition) of the pathway (Figure 7A). Similar to *STAT1* mRNAs, *IFIT1* and *MX1* transcripts were rapidly degraded in the NAT10 KO cells, with half-lives of 0.9- and 0.8 hours respectively, compared to 3.4- and 4.8 hours in WT cells (Figure 8B, 8C). To exclude the possibility that global mRNA degradation increased in the NAT10 KO cells, we also examined *TP53*, a stress-induced transcription not involved in the innate immune response pathway yet affected during ZIKV infection (63–65). Both p53 mRNA and protein were previously reported to be increased during ZIKV infection and contribute to apoptosis in human neuronal progenitor cells (65). We found that *TP53* mRNA stability was similar in both WT and NAT10 KO cells (Figure 8D). In summary, these data show that NAT10 affects the stability of *STAT1*, *IFIT1* and *MX1*, three cellular transcripts involved in an antiviral response. Overall, these data indicate that the earlier and higher expression of ZIKV protein and RNA leading to increased virus replication, are directed in part by either NAT10 or because of ac^4^C modifications on RNA, affecting the stability of select innate immune response genes.

**Figure 8:**
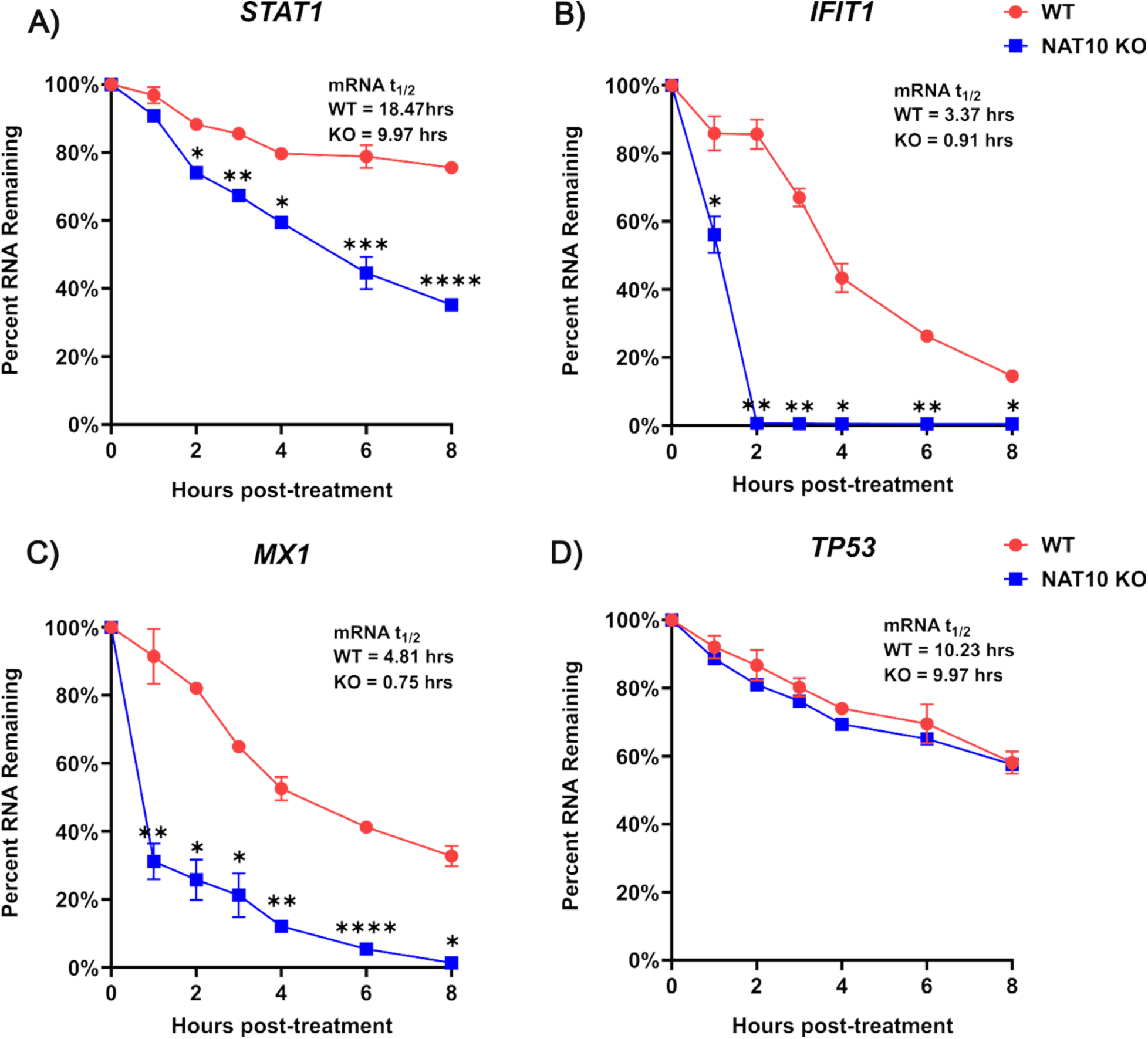
*STAT1*, *IFIT1* and *MX1* mRNAs decay more rapidly in the absence of NAT10. A549 WT and NAT10 KO cells were infected with ZIKV at a MOI of 5 pfu/cell. Sixteen hours post-infection, cells were incubated with new media containing either DMSO or 4μM Actinomycin D. At the time of harvest (0-, 1-, 2-, 3-, 4-, 6- and 8-hours post-treatment), cells were lysed in TRIzol and RNA isolated. *STAT1*, *IFIT1, MX1, and TP53* RNA levels quantified relative to *ACTB* mRNA by RT-qPCR. The mRNA decay data are from three independent experiments. RNA levels were normalized to the 0-hour timepoint and converted into a percentage. A 2-way ANOVA was used to determine statistical significance. Error bars show ± SD. The data at each time from each RNA analyzed were determined to be statistically significant. For simplicity, only *P<0.05, **P<0.01 and ***P<0.001 are annotated on the graph. Unannotated data show no significance.

## DISCUSSION

Although RNA modifications have been known for decades, the functional consequences of these chemical moieties have largely remained poorly understood. Recent advances in detection technologies, however, are beginning to uncover the role of these chemical marks play in regulation of RNA function (66). Notably, significant progress has been made towards defining the biological function of N6-methyladenosine, pseudouridine, 5-methylcytosine, 2’-O-methylations and N4-acetylcytosine (ac^4^C) in both cellular and viral RNAs (22, 41–43, 46–49, 62, 67–73). In an earlier study, we determined that the levels of the ac^4^C modification decreased during ZIKV infection, and that the modification was present on the viral RNA (22). To explore the functional significance of ac^4^C in this context we investigated the role of NAT10, the writer enzyme, during ZIKV infection. Our findings revealed that NAT10 regulated the stability of innate immune response genes that function to suppress ZIKV gene expression.

Ac^4^C has crucial roles in a variety of cellular and viral RNAs (42–44, 46–49, 62, 67, 69, 73). Using crosslinking and immunoprecipitation (CLIP) approaches with either an antibody recognizing the ac^4^C modification and/or NAT 10 protein, putative ac^4^C marks were reported on HIV-1 and EV71 viral RNAs (47, 48, 73). In HIV-1, the ac^4^C sites were distributed along the viral RNA, with higher acetylation of cytidine in the *pol* and *env* polyprotein ORFs, and long terminal repeat regions, depending on the CLIP approach used (47). Using NAT10 KO cells, Tsai and colleagues showed a decrease in HIV-1 protein and RNA levels resulting from increased turnover of the HIV-1 RNA (47). An orthogonal approach of mutating putative ac^4^C sites within the HIV-1 *env* polyprotein ORF, similarly showed some decrease in viral protein, RNA and titers, although the functional consequence of such mutagenesis was not investigated (47). In contrast, we found that in NAT10 KO cells, ZIKV protein and RNA increased, and this increase was in part the result of earlier viral gene expression (Figure 3), and increased ZIKV replication (Figure 4 and Figure 5). Like the findings in HIV-1 (47) we observed increased turnover of ZIKV RNA (Figure 4B, Figure 4D and Figure 5B), as indicated by the decrease in *Renilla* expression from the subgenomic reporter replicon. With the increased decay of ZIKV RNA we initially posited that more ZIKV sfRNAs would accumulate, which would alter the immune response. Surprisingly, however, by Northern blot we observed no difference in the abundance of sfRNAs in WT and NAT10 KO cells (Figure 5C). Notably, Arango et al., (2018) showed that XRN1 decay of the RNA was not affected by the absence or presence of ac^4^C on mRNAs (42). The disconnect between ZIKV RNA turnover and sfRNA abundance hints at alternative decay mechanisms that may act independently of XRN1.

The presence of ac^4^C marks within rRNA (43, 44, 46), tRNA (46) and mRNA (42) support a role for NAT10 and these RNA modifications in translation. Indeed, Arango and colleagues showed that ac^4^C marks in the wobble cytidine position in an mRNA transcript stimulated translation likely by promoting interactions with the cognate tRNA (42). Cytidines within the EV71 internal ribosome entry site (IRES) are N4-acetylated, and depletion of NAT10 and mutation of the specific modified cytidine nucleotides decreased EV71 protein, RNA and titers (48). The resulting decrease in translation efficiency was shown to be a result of the reduced binding of the IRES transacting factor PCBP1, and hence the binding of the ribosome to the EV71 IRES (48). The consequence of reduced translation of EV71 also increased turnover of the viral RNA (48). Given the reported role for NAT10 on translation, and the increase in ZIKV protein levels, we expected that ZIKV translation might also be modulated by NAT10. Using the subgenomic *Renilla* luciferase reporter replicons we, however, did not find a consistent effect on translation (Figure 4 and Figure 5). For example, at early time points (4- and 8-hours) post-transfection we observed no difference in translation of the replication competent reporter replicon RNA. In contrast however, translation of the replication incompetent replicon RNA was higher in the NAT10 KO cells compared to WT cells (Figure 4 and Figure 5). Consistent between both replicon RNAs however was an effect on replication, which in the absence of NAT10 was increased at earlier time points (Figure 4 and Figure 5). In addition to promoting translation, ac^4^C marks on EV71 RNA increased the binding of the 3D RNA-dependent RNA polymerase (RdRp) to the viral RNA, which promoted replication of the positive-sense RNA but not the negative-sense RNA (48). Curiously, NAT10 was previously reported to bind both Dengue virus serotype 2 (strain 16882) and ZIKV NS5 RdRp protein (74). While the consequence of this interaction was not investigated (74), it is possible that NAT10 regulates ZIKV replication by modulating the subcellular localization and/or activity of NS5. This interaction and novel regulation of flavivirus NS5 proteins by NAT10 is one focus of future studies.

Despite the increase in ZIKV protein and RNA, at 48-hours post infection in the HeLa and A549 NAT10 KO cells we observed no difference in viral titers (Figure 1, Figure 6C and Figure 7C). These data contrast with the studies with HIV-1, EV71 and SINV (47, 48, 62), where viral gene expression and titers were coupled. When we examined viral titers at 24 hours post-infection, we observed that the increase in virus titers coincided with the increased viral protein and RNA levels (Figure 6C and Figure 7C). We speculated that NAT10, directly or indirectly, might affect the assembly and trafficking of ZIKV. However, when we measured virus titers from cells (intracellular) versus those released into the media (extracellular) we observed no significant difference (Figure 3). Interestingly, NAT10 was previously reported to interact with the capsid protein from both Dengue virus and ZIKV (74), which might contribute to the production of infectious particles. Additional studies are however needed to determine if NAT10 has a post-replication function (e.g., in virus assembly and/or release) that may explain the apparent uncoupling between viral gene expression and production of infectious virions.

The effect of NAT10 KO and absence of ac^4^C in cellular mRNA was previously shown to affect RNA turnover and translation (42, 67). 5-Bromouridine labeling and isolation of cellular RNAs and analysis by next generation RNA sequencing revealed that cellular mRNAs modified with ac^4^C degraded more rapidly than unmodified mRNAs (42). Rather than labeling newly synthesized RNA, we inhibited cellular RNA polymerase II transcription with actinomycin D, measured relative RNA abundance by RT-qPCR and then calculated the mRNA half-life from the resulting decay curves (Figure 8). Our data are consistent with the increased turnover of innate immune response mRNAs reported by Arango and colleagues (42). Notably, in our analysis, we observed the rapid decay of *STAT1*, *IFIT1* and *MX1* mRNAs in the NAT10 KO cells compared to WT cells during ZIKV infection (Figure 8). We determined that the change in stability was specific to the analyzed mRNAs in the NAT10 KO cells, as the half-life of *TP53*, a transcript that was previously not reported to be modified with ac^4^C (42), was similar in both ZIKV infected WT and NAT10 KO (Figure 8). These data support a role of NAT10 in the innate immune response pathway. Indeed, the reduced amount of *IFNB* (Figure 6D), and the decreased mRNA stability of *STAT* mRNA and reduced protein expression (Figure 7), likely attenuated the signaling within the JAK/STAT pathway. We validated this by treating cells with Ruxolitinib, a JAK1/2 inhibitor. Ruxolitinib treatment inhibited the expression of IFIT1 and MX1 (Figure 7) and resulted in comparable expression of ZIKV protein and RNA, and production of infectious virions in both WT and KO cells (Figure 7). In addition, the attenuated signal cascade from reduced STAT1 levels was likely further compromised by the decreased stability of *IFIT1* and *MX1*.

During SINV infection, the abundance of NAT10 and ac^4^C modifications increased, and NAT10 was reported to affect SINV infection (62). Dang and colleagues showed that when NAT10 was depleted with shRNAs, or mutations were introduced into NAT10 to disrupt the RNA helicase and N-acetyltransferase activities, SINV protein, RNA and titers decreased (62). RNA-seq analysis showed the up- and downregulation of different cellular transcripts. The authors identified that lymphocyte antigen six family member E (*LY6E*) mRNA was downregulated (62). Indeed, *LY6E* was previously shown to be ac^4^C modified (42). Like our study, this ISG was rapidly degraded in NAT10 depleted cells (62). Similarly, during KSHV infection, the interferon-inducible protein 16 (*IFI16*) mRNA was reported to be acetylated by NAT10 which promoted mRNA stability and translation leading to activation the inflammasome (73). NAT10 was also previously described to regulate the stability of UNC-52-like kinase 1 (*ULK1*) mRNA in neutrophils, which when decreased induced the activation of the STING-IRF3 signaling pathway and NLRP3 inflammasome (75). An immune study demonstrated that during T-cell activation the levels of NAT10 increased (72, 76). Furthermore, in the absence of NAT10, the acetylation of RNAs was diminished and T-cells proliferation reduced, which in turn affected the antiviral response to lymphocytic choriomeningitis virus infection (72). These reports, combined with our data, suggest that NAT10 may act as a selective regulator of mRNA stability within the innate immune response, a function that to date has not been fully appreciated.

In this study we extend our understanding of and show a role for NAT10, the ac^4^C writer enzyme on the stability of three components within the innate immune response pathway during ZIKV infection. We used both HeLa and A549 WT and NAT10 KO cells. In the NAT10 KO cells we found that ZIKV protein and RNA levels increased (Figure 1 and Figure 6), but viral titers were only affected at early time points (Figure 6). These data are consistent with an earlier report by Serman and colleagues who investigated the acetylation of ZIKV proteins by cellular lysine acetyltransferases (77). These investigators reported that depletion of NAT10 increased the amount of ZIKV E RNA (77). We found that the increase in viral gene expression was a result of altered infection kinetics (Figure 1 and Figure 3), which increased ZIKV replication (Figure 4 and Figure 5A). In the NAT10 KO cells, the ZIKV RNA appeared to be turned over rapidly (Figure 4 and Figure 5B). We initially posited that with the increased ZIKV RNA turnover, the levels of sfRNA would be increased, which through increased sfRNA levels and sequestering of innate immune factors (78, 79) would contribute to promoting ZIKV infection. However, we found no difference in the accumulation of sfRNA between WT and NAT10 KO cells (Figure 5C). Rather, we determined that in cells devoid of NAT10, the stability of select innate immune response factors decreased (Figure 7 and Figure 8), thus attenuating this critical antiviral pathway and shifting the infection kinetics to earlier time points to increase ZIKV gene expression (Figure 9). Overall, our study expands on the role of RNA modifications on viral infections and provides additional insight into how the viruses might modulate the innate immune response by restricting NAT10 expression and ac^4^C modifications on cellular RNAs.

**Figure 9:**
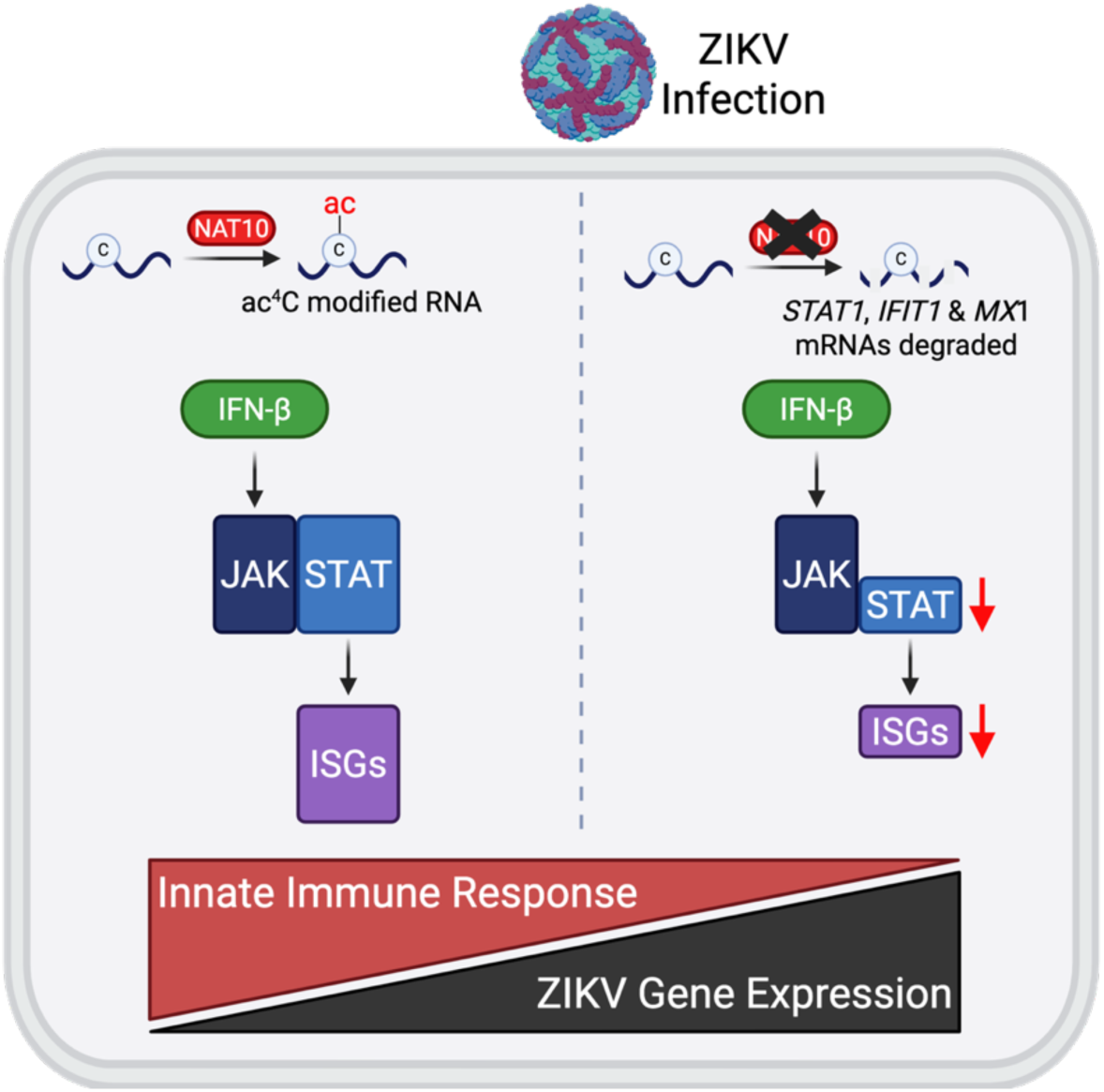
NAT10 promotes mRNA stability of select immune response factors to modulate ZIKV infection. NAT10 is a writer enzyme that deposits an acetyl group on cytidine to form N4-acetylcytidine (ac^4^C). In response to viral infection, the host cell initiates a robust innate immune response through a primary signaling cascade to induce the expression of type I interferons (e.g. interferon β, IFN-β). The release of IFN-β in turn activates the secondary JAK/STAT signaling pathway to induce expression of interferon stimulated gene (ISGs), such as *IFIT1* and *MX1*, which robustly act to restrict ZIKV infection. However, when NAT10 is absent several mRNAs associated with the immune response pathway (e.g., *STAT1*, *IFIT1*, and *MX1)* are rapidly degraded resulting in decreased protein levels and an attenuated innate immune response. The reduced cellular response shifts the viral kinetics to earlier time points and increases ZIKV gene expression.

## MATERIALS AND METHODS

### Cell Lines and Zika Virus

HeLa (human cervical epithelial adenocarcinoma) wild type (WT) and NAT10 knockout (KO) cell lines (a kind gift from Dr. Jordan Meier, NCI, (42)) were maintained in Dulbecco’s minimal essential medium (DMEM; Gibco, #11995–065) supplemented with 10% fetal bovine serum (FBS; Seradigm, #97068–085), 2 mM L-glutamine (Gibco, #25030081), and 1 mM sodium pyruvate (Gibco, #11360070). A549 (Human lung epithelial adenocarcinoma, ATCC CCL-185) WT and NAT10 KO cell lines were maintained in DMEM (Gibco, #11995–065) supplemented with 10% FBS (Seradigm, #97068–085), 2 mM L-glutamine (Gibco, #25030081), and 1 mM sodium pyruvate (Gibco, #11360070). Vero cells (ATCC CRL-81) were cultured in DMEM (Gibco, #11995–065) supplemented with 10% FBS (Seradigm, #97068–085), 1% penicillin and streptomycin (Gibco, #15140163), and 10 mM HEPES (Gibco, #15630080). HEK293 FT cells (Life Technologies) were grown in DMEM with 10% FBS (Seradigm, #97068–085), 10 mM nonessential amino acids (NEAA; Gibco, #11140076) and 2 mM L-glutamine (Gibco, #25030081). All cell lines were cultured at 37°C with 5% CO_2_ in a water-jacketed incubator. C6/36 (ATCC CRL-1660) were grown in EMEM (Thermo Scientific) with 10% FBS (Seradigm, #97068–085), 1% penicillin and streptomycin (Gibco, #15140163), 1 mM sodium pyruvate (Gibco, #11360070), and 1% Amphotericin B (Gibco, #15290026), at 28°C with 5% CO_2_ in a water-jacketed incubator.

The ZIKV (Uganda MR766) full length infectious clone (pCDNA6.2 ATCCMR766 Intron3127 HDVr) was a generous gift from Dr. Matthew Evans (Icahn School of Medicine at Mount Sinai) (80). To generate new infectious virus stocks, Vero cells seeded in a 6-well plate at 3×10^5^ cells/well were transfected with 1 μg of the infectious clone plasmid using Lipofectamine 3000, per the manufacturer’s method. The media was collected five days post-transfection and aliquoted. The initial stock of virus was amplified once on Vero cells and harvested 48 hours post-infection. Final stocks of virus used throughout this manuscript were collected after two additional amplifications on C6/36 cells each harvested 5 days post-infection. Virus stocks were collected and supplemented with 20% FBS, aliquoted, and stored at -80°C. We validated the infection by RT-qPCR and measured viral titers by plaque assay.

### ZIKV Infection

Twenty-four hours prior to the infection, cells were seeded at 1x 10^6^ cells in a 10 cm cell culture dish. On the day of the infection, a mock plate of seeded cells was trypsinized and counted to determine the virus concentration to infect the cells at. An appropriate aliquot of viral stock was thawed at room temperature, diluted in PBS to a final volume of 1 mL and added to cells. All cells were infected at a multiplicity of infection (MOI) of 5 plaque forming units (pfu)/cell. For mock-infected plates, 1 mL of PBS was added. Cells were rocked and then incubated at 37°C for 1 hour. Hereafter, 9 mL of media was added to each dish and the cells were further incubated at 37°C. At the respective time point, cells were harvested for analysis of protein, RNA and ZIKV titers. For viral titers, duplicate 500 μL aliquots of media were stored at -80°C.

### Harvest of Infected and Chemically Treated Cells

Mock- and virus-infected and chemically treated cells were harvested as follows; first, media was collected in aliquots for downstream analysis of viral titers. Cells were gently washed twice with 2 mL cold PBS (Gibco, #14190250) and the PBS aspirated. A volume of 1 mL cold PBS (Gibco, #14190250) was then added to the plates, cells were scraped off the plate using a cell lifter, and the cell suspension was mixed thoroughly. Equal volumes of 500 µL were aliquoted into two separate tubes. Cell suspensions were centrifuged at 14,000 rpm for 30 seconds to pellet the cells. The PBS supernatant was aspirated and cells in one tube were prepared for protein analysis, while the other tube was prepared for RNA analysis.

### CRISPR-Cas9 Gene Editing of NAT10 and THUMPD1

We used the CRISPR-Cas9 system to generate A549 NAT10 KO cells and HeLa THUMPD1 KO cells. Specifically, a gRNA sequence (5’-GTGAGTTCATGGTCCGTAGG-3’; Genscript) (42) targeting NAT10 and a gRNA sequence (5’-CCACCGGATGTCCTCTGCGC-3’; Genscript) targeting THUMPD1 was cloned into pLentiCRISPRv2 plasmid. HEK 293 FT cells were co-transfected with pLentiCRISPRv2-NAT10 CRISPR gRNA or with pLentiCRISPRv2-THUMPD1 CRISPR gRNA and pMD2.G (Addgene, #12259) and psPAX2 (Addgene, #12260) packaging plasmids using JetOptimus DNA transfection reagent (Polyplus, #101000025) according to the manufacturer’s protocol. Media containing lentivirus at 24- and 48-hours post-transfection was collected, pooled, filtered through a 0.45 mm pore filter and then used to transduce A549 or HeLa cells in the presence of 6 µg/mL polybrene (Sigma-Aldrich, TR1003). After 48 hours of incubation at 37°C, the media on the A549 or HeLa cells was removed and replaced with media containing puromycin (1 µg/mL; InvivoGen, #ant-pr-1), and selection was carried out for 4 days by which time all A549 or HeLa WT control cells were killed by the antibiotic. The cell suspension was diluted, and individual cells seeded into a 96-well plate and then incubated at 37°C. Following expansion, clones were screened for NAT10 or THUMPD1 expression by Western blotting and RT-qPCR. CRISPR-Cas9 gene editing of NAT10 or THUMPD1 was validated by sequencing. Specifically, DNA was isolated from KO cells using DNAzol (Invitrogen, #10503027) reagent. PCR was subsequently carried out with forward and reverse primers (5’-TGGCTTTGTGCTCTGAAGTC-3’ and 5’-GCTCTTAGCCCAGAGGCTGT-3’) designed to exon 5 for NAT10 while PCR was subsequently carried out with primers (forward: 5’-TCGCGCCGGCGATAGAC-3’ and reverse: 5’-CGCGCATGCCCATACCA-3’) designed to exon 2 for THUMPD1. The PCR products were cloned into pCR2.1 vector via TOPO cloning system (Invitrogen # ) and the sequence analyzed by Sanger sequencing.

### Chemical Treatments

Ruxolitinib, a selective inhibitor of JAK1 and JAK2 (60, 61), was reconstituted in DMSO to a stock concentration of 10 mM. A549 WT and NAT10 KO cells were first mock- and ZIKV-infected at a moi of 5 pfu/cell for 1 hour at 37°C. DMSO (control) and Ruxolitinib (Selleck Chem, #S1378) at 30 nM final concentration was added to the replaced media and maintained in the culture media for the duration of infection. Cells were harvested as described above. Actinomycin D, an RNA polymerase II inhibitor (Sigma, A1410-2MG) (81), was reconstituted in DMSO to a stock concentration of 2mM. A549 WT and NAT10 KO cells were either mock-infected or infected with ZIKV (MOI = 5 pfu/cell). At 16 hours post-infection, the media was removed and replaced with new media containing either 4 μM Actinomycin D or the equivalent volume of DMSO. DMSO or Actinomydin D-treated cells were lysed in TRIzol at the time the media was replaced (0 hours) and at 1-, 2-, 3-, 4-, 6-, and 8-hours post-treatment.

### siRNA Transfections and Plasmid Transfections

The siRNA targeting NAT10 was purchased from Invitrogen (ID:123432). We used siGL2 targeting *Guassia* Luciferase (sense-strand 5’-CGUACGCGGAAUACUUCGAUU-3’ and antisense strand 5’-UCGAAGUAUUCCGCGUACGUU-3’) as a control siRNA. The siRNA duplex was prepared as previously described (82). Huh7 WT cells were seeded in a 6 cm cell culture dish at a concentration of 7×10^5^ cells. After 24 hours, the cells were transfected with 100 nM siRNAs using Lipofectamine 3000 per the manufacturer’s protocol. The following day the cells were reseeded into a 10 cm cell culture dish and 48 hours post-transfection infected with ZIKV at an MOI of 5 pfu/cell. Cells were harvested at 48 hours post-infection.

We used JetOPTIMUS transfection reagent (Polyplus, # 101000006) according to the manufacturer’s instructions to introduce the 3xFlag-NAT10 plasmid (a kind gift from Dr. Jordan Meier, NCI, (42, 83)) and the 3xFlag-Bacterial Alkaline phosphatase (BAP; Sigma, #E7533) into HeLa NAT10 KO cells. Cells were seeded at 1×10^6^ in 10 cm cell culture dishes and after 24 hours transfected with transfection reagent alone (vehicle control) or 5 μg of plasmid DNA. Cells were infected with ZIKV (MOI=5 pfu/cell) and then total protein, RNA, and media harvested at 48 hours post-infection.

### Western Blot Analysis

To analyze ZIKV and cellular proteins, cells were lysed with RIPA buffer (100 mM Tris-HCl pH 7.4, 0.1% sodium dodecyl sulphate (SDS), 1% Triton X-100, 1% deoxycholic acid, 150 mM NaCl) containing protease inhibitors and phosphatase inhibitors (EDTA-free; ThermoScientific, #A32965, #A32957) and incubated on ice for 20 minutes. Clarified supernatant was collected after the cell lysates were centrifuged at 4°C for 20 minutes at 14,000 rpm. The concentration of proteins in the lysate was quantified using the DC protein assay kit (Bio-Rad, # 5000112). Twenty-five micrograms of proteins were separated in a 10% SDS-polyacrylamide (PAGE) gel at 100 V for 2 hours, and then transferred on to polyvinylidene difluoride membrane (Millipore Sigma, # IPVH08100) at 30V overnight or 100V for 1 hour at 4°C. The blots were activated in absolute methanol and stained with PonceauS (Sigma, # P7170-1L) to determine transfer efficiency. Hereafter, blots were washed in PBS buffer with 0.1% Tween (PBS-T) and blocked in 5% milk in PBS-T for 1 hour at room temperature. The blots were incubated with primary antibodies diluted in either 5% milk or 5% BSA in PBS-T for 1 hour at room temperature or overnight at 4°C. The next day the blots were washed thrice for 10-minute PBS-T washes and then incubated in secondary antibodies in 5% milk in PBS-T for 1 hour at room temperature.

The blots were washed three times in PBS-T and the proteins were visualized using Clarity Western ECL blotting substrate (Bio-Rad) or SuperSignal West Femto (ThermoScientific). The primary and secondary antibodies used are listed in Table 1.

**Table 1.**
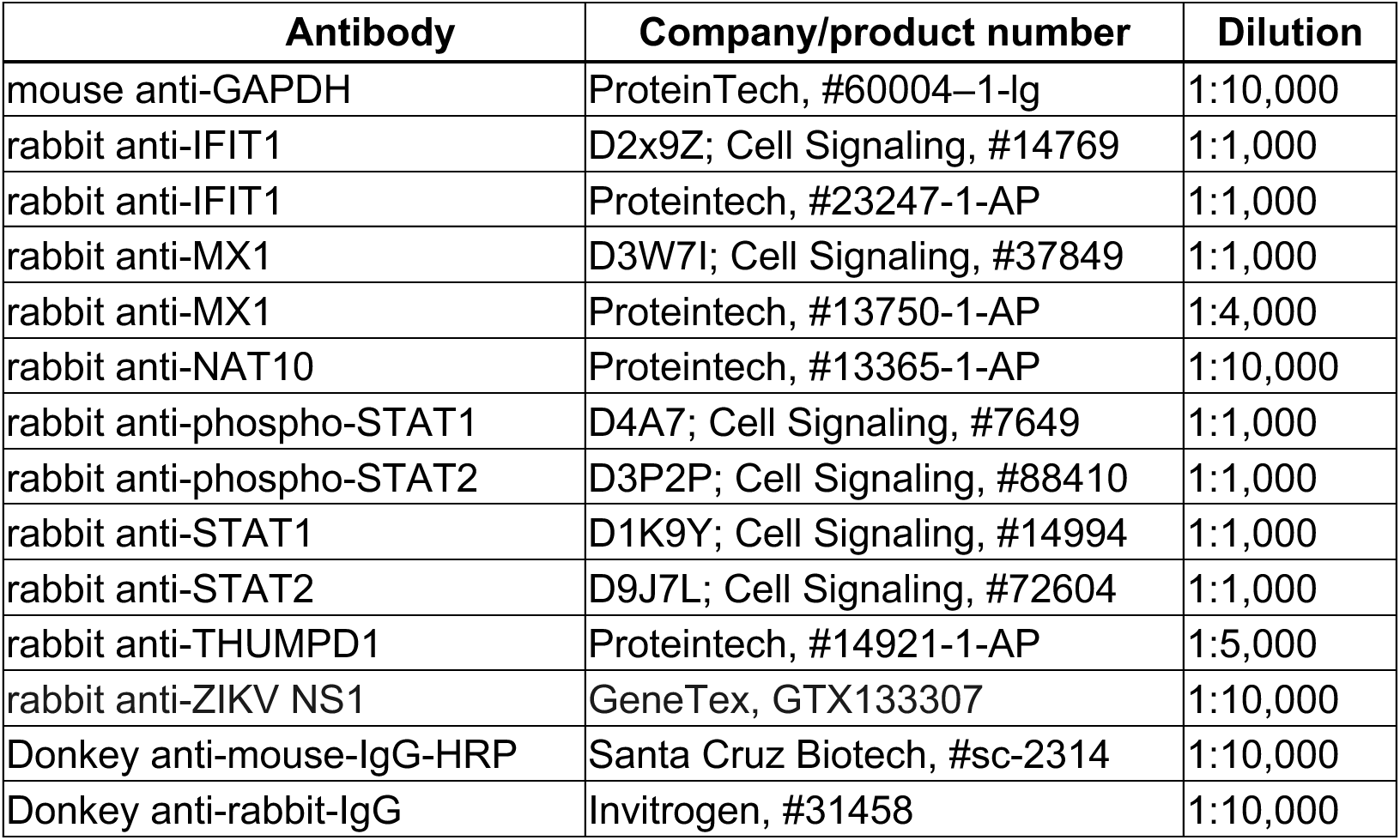

### RT-qPCR Analysis

Total RNA was isolated from cells using TRIzol reagent (Invitrogen, #15596026) and the RNA Clean and Concentrator kit (Zymo Research, #R1018). The RNA was DNase-treated using the TURBO DNA-free kit (Invitrogen, #AM1907) and reverse transcribed using the High-Capacity cDNA Reverse Transcription reagents (Applied Biosystems, #4368813). The resulting cDNA was used for qPCR analysis with iTaq Universal SYBR Green Supermix reagents (Biorad, #1725124) and CFX384 Touch Real-Time PCR system (Biorad). RT-qPCR data shown are from at least three independent experiments, with each sample assayed in three technical replicates.

Table 2 lists the primers used in our analysis.

**Table 2.**
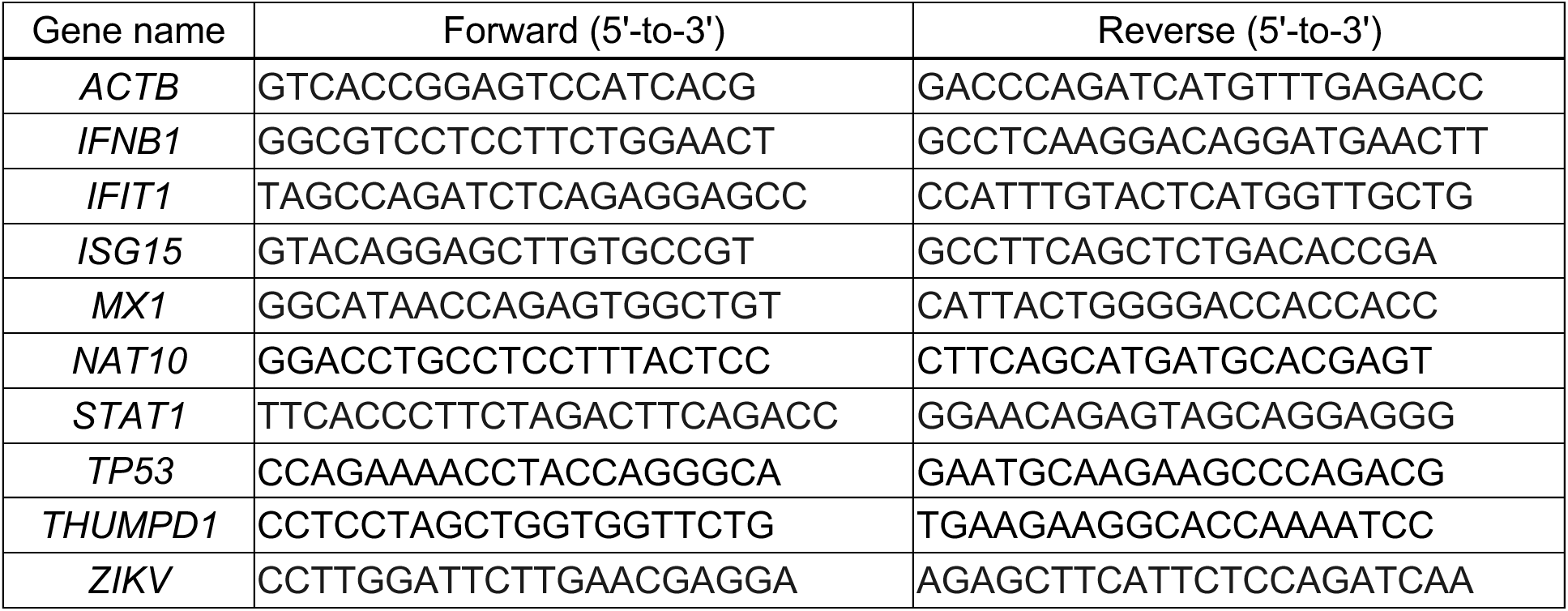

### Plaque Assays

Vero cells were seeded in six-well plates at a density of 7×10^5^ cells/well and incubated at 37°C with 5% CO2 overnight. The following day, 10-fold serial dilutions from 10^−1^ to 10^−6^ of media from infections were prepared in 1x PBS (Gibco, #14190250). The media on Vero cells seeded the previous day were aspirated, 300 µL of 1x PBS was added to the mock well, and 300 µL of each virus dilution was added to the remaining wells. The cells were incubated at 37°C with 5% CO2 for 1 hour, gently rocking every 15 minutes. After incubation, the PBS or virus dilution in PBS was aspirated and 3 mL of overlay consisting of 1:1 2x DMEM [DMEM high glucose, no sodium bicarbonate buffer powder (Gibco # 12–100-046) in 500 mL of RNase-free water, 84 mM of sodium bicarbonate, 10% FBS, and 2% penicillin and streptomycin, at pH 7.4] and 1.2% avicel (FMC, #CL-611) was added to each well, and the plates were incubated at 37°C with 5% CO2. Five days post-infection, the overlay was aspirated, cells were fixed with 1 mL of 7.4% formaldehyde (Fischer Scientific, #F79-500) for 10 minutes at room temperature and rinsed with water. Plaques were visualized using 1% crystal violet (Sigma, #C3886) in 20% methanol. Viral titers were determined from duplicate viral dilutions and three biological replicates.

To determine the titer of ZIKV infectious particles inside the cells, we trypsinized and collected cells in a 1.7mL microcentrifuge tube. Cells were counted using a hemocytometer and 1×10^5^ of cells from each sample was aliquoted. Collected cells were centrifuged at 10,000 rpm for 30 seconds and then resuspended in 150 µL of PBS and stored at -80°C. Fifty µL of thawed cell lysate was used in subsequent plaque assays being used to create each serial dilution. Plaque assays to determine intracellular virus was performed in technical duplicate for each biological replicate of the assay.

### *In Vitro* Transcription and Transfection of sgRLuc-ZIKV and Luciferase Analysis

*Renilla* luciferase assays to investigate translation and replication in HeLa and A549 WT and NAT10 KO were performed using the previously described replication competent and incompetent synZIKV-R2A replicon (53). Plasmid DNA was transformed in NEB Turbo (#C2984H) and isolated using the Omega Bio-Tek E.N.Z.A Plasmid DNA Mini Kit I (#D6942-00S). The reporter DNA was linearized using XhoI (NEB, #R0146S), purified by phenol-chloroform extraction and then precipitated using 0.5 volume of 5M ammonium acetate (G-Biosciences, #R012), 1 µL 1 mg/mL glycogen (Invitrogen, #AM9510), and 2.5 volumes of absolute ethanol at -20°C overnight. The linearized template DNA was pelleted by centrifugation at 14,000 rpm for 20 minutes, the pellet washed with 70% ethanol and then resuspended in 10 µL of RNAse-free water. Hereafter, the template DNA was *in vitro-* transcribed using the HiScribe T7 High Yield RNA Synthesis Kit (NEB #E2040S) and 5’-end capped using the 3’-O-Me-m7G(5’)ppp(5’)G RNA Cap Structure Analog (NEB #S1411L) as recommended by the manufacturer. Reactions were incubated at 37°C for two hours. We validated and visualized the RNA production by electrophoresis in a 0.8% native agarose gel with Gel Red (Biotinium, #41003). The remaining *in vitro* transcription reaction was treated with 0.1 volume DNAseI (NEB #M0303) for 15 minutes. The RNA was purified by phenol chloroform extraction and precipitated overnight at -80°C. Prior to cell transfection, the RNA was pelleted and resuspended in RNAse-free water. Both A549 and HeLa cells were seeded in 24 well plates at 7×10^4^ cells/well 24 hours prior to transfection. After 24 hours, the cells were transfected with 1 µg of RNA/well using Lipofectamine 3000 per the manufacturer’s protocol.

At each time point, the media was aspirated, cells were washed once with PBS and then lysed in 350 µL of 1X *Renilla* luciferase lysis buffer (*Renilla* Luciferase Assay System, Promega #E2810) per the manufacturer’s instructions. Cell lysates were stored at -20°C prior to analysis. *Renilla* luciferase activity in technical duplicate or triplicate was measured using a Promega GloMax luminometer. Specifically, 20 µL of cell lysate in a white 96-well plate was assayed with 50 µL of 1:100 diluted *Renilla* luciferase reaction solution per well.

### Statistical Analysis

The data shown are from at least three independent experiments. Data was analyzed using Prism 9.4.1 software (GraphPad, La Jolla, CA, USA) to establish statistical significance. We performed two-tailed students t-test for two groups and two-way ANOVA for multiple group comparisons.

## ACKNOWLEDGEMENTS

This work was supported by grants from National Institutes of Health to CTP (R01GM123050 and R21AI178672). The research in this manuscript is solely the responsibility of the authors and does not necessarily represent the official views of the NIH. We gratefully acknowledge Dr. Jordan Meier (NCI) for advice as we embarked on this research and for generously sharing the HeLa WT and NAT10 KO cells and 3xFlag-tagged NAT10 plasmid with us. We also thank Dr. Matthew Evans (Icahn School of Medicine at Mount Sinai) and Dr. Ralf Bartenschlager (Heidelberg University) for respectively sharing the ZIKV infectious clone (pCDNA6.2 ATCCMR766 Intron3127 HDVr) and subgenomic ZIKV *Renilla* luciferase reporter replicon (synZIKV-sgR2A replicons) with us. Last, we appreciate the NYSDOH Wadsworth Arbovirus laboratory and CDC for permission to use ZIKV PRVABC59. Last, we thank the Pager lab for their thoughtful comments and suggestions on this manuscript.

## REFERENCES

1. Dick GWA. 1952. Zika Virus (I). Isolations and serological specificity. Trans R Soc Trop Med Hyg 46:509–520.

2. Dick GWA. 1952. Zika virus (II). Pathogenicity and physical properties. Trans R Soc Trop Med Hyg 46:521–534.

3. Duffy MR, Chen T-H, Hancock WT, Powers AM, Kool JL, Lanciotti RS, Pretrick M, Marfel M, Holzbauer S, Dubray C, Guillaumot L, Griggs A, Bel M, Lambert AJ, Laven J, Kosoy O, Panella A, Biggerstaff BJ, Fischer M, Hayes EB. 2009. Zika Virus Outbreak on Yap Island, Federated States of Micronesia. New England Journal of Medicine 360:2536–2543.

4. Lazear HM, Diamond MS. 2016. Zika Virus: New Clinical Syndromes and Its Emergence in the Western Hemisphere. J Virol 90:4864–4875.

5. Gubler DJ, Vasilakis N, Musso D. 2017. History and Emergence of Zika Virus. J Infect Dis 216:S860–S867.

6. Musso D, Gubler DJ. 2016. Zika virus. Clin Microbiol Rev 29:487–524.

7. Musso D, Nilles EJ, Cao-Lormeau VM. 2014. Rapid spread of emerging Zika virus in the Pacific area. Clinical Microbiology and Infection 20:O595–O596.

8. Musso D, Roche C, Robin E, Nhan T, Teissier A, Cao-Lormeau VM. 2015. Potential sexual transmission of zika virus. Emerg Infect Dis 21:359–361.

9. Grant R, Fléchelles O, Tressières B, Dialo M, Elenga N, Mediamolle N, Mallard A, Hebert JC, Lachaume N, Couchy E, Hoen B, Fontanet A. 2021. In utero Zika virus exposure and neurodevelopment at 24 months in toddlers normocephalic at birth: a cohort study. BMC Med 19:12.

10. Marbán-Castro E, Goncé A, Fumadó V, Romero-Acevedo L, Bardají A. 2021. Zika virus infection in pregnant women and their children: A review. European Journal of Obstetrics and Gynecology and Reproductive Biology 265:162–168.

11. Styczynski AR, Malta JMAS, Krow-Lucal ER, Percio J, Nóbrega ME, Vargas A, Lanzieri TM, Leite PL, Staples JE, Fischer MX, Powers AM, Chang GJJ, Burns PL, Borland EM, Ledermann JP, Mossel EC, Schonberger LB, Belay EB, Salinas JL, Badaro RD, Sejvar JJ, Coelho GE. 2017. Increased rates of Guillain-Barré syndrome associated with Zika virus outbreak in the Salvador metropolitan area, Brazil. PLoS Negl Trop Dis 11: e0005869.

12. Cao-Lormeau VM, Blake A, Mons S, Lastère S, Roche C, Vanhomwegen J, Dub T, Baudouin L, Teissier A, Larre P, Vial AL, Decam C, Choumet V, Halstead SK, Willison HJ, Musset L, Manuguerra JC, Despres P, Fournier E, Mallet HP, Musso D, Fontanet A, Neil J, Ghawché F. 2016. Guillain-Barré Syndrome outbreak associated with Zika virus infection in French Polynesia: A case-control study. The Lancet 387:1531–1539.

13. França GVA, Schuler-Faccini L, Oliveira WK, Henriques CMP, Carmo EH, Pedi VD, Nunes ML, Castro MC, Serruya S, Silveira MF, Barros FC, Victora CG. 2016. Congenital Zika virus syndrome in Brazil: a case series of the first 1501 livebirths with complete investigation. The Lancet 388:891–897.

14. Ryan SJ, Carlson CJ, Tesla B, Bonds MH, Ngonghala CN, Mordecai EA, Johnson LR, Murdock CC. 2021. Warming temperatures could expose more than 1.3 billion new people to Zika virus risk by 2050. Glob Chang Biol 27:84–93.

15. Wang L, Jia Q, Zhu G, Ou G, Tang T. 2024. Transmission dynamics of Zika virus with multiple infection routes and a case study in Brazil. Sci Rep 14:7424.

16. Ryan SJ, Carlson CJ, Mordecai EA, Johnson LR. 2019. Global expansion and redistribution of Aedes-borne virus transmission risk with climate change. PLoS Negl Trop Dis 13:e0007213.

17. Guirakhoo F, Bolin RA, Roehrig JT. 1992. The Murray Valley encephalitis virus prM protein confers acid resistance to virus particles and alters the expression of epitopes within the R2 domain of E glycoprotein. Virology 191:921–931.

18. Hasan SS, Sevvana M, Kuhn RJ, Rossmann MG. 2018. Structural biology of Zika virus and other flaviviruses. Nature Structural & Molecular Biology 25:13–20.

19. Munjal A, Khandia R, Dhama K, Sachan S, Karthik K, Tiwari R, Malik YS, Kumar D, Singh RK, Iqbal HMN, Joshi SK. 2017. Advances in developing therapies to combat zika virus: Current knowledge and future perspectives. Front Microbiol 8:287094.

20. O’Connor MA, Tisoncik-Go J, Lewis TB, Miller CJ, Bratt D, Moats CR, Edlefsen PT, Smedley J, Klatt NR, Gale M, Fuller DH. 2018. Early cellular innate immune responses drive Zika viral persistence and tissue tropism in pigtail macaques. Nature Communications 9:3371.

21. Ye Q, Liu ZY, Han JF, Jiang T, Li XF, Qin CF. 2016. Genomic characterization and phylogenetic analysis of Zika virus circulating in the Americas. Infection, Genetics and Evolution 43:43–49.

22. Mcintyre W, Netzband R, Bonenfant G, Biegel JM, Miller C, Fuchs G, Henderson E, Arra M, Canki M, Fabris D, Pager CT. 2018. Positive-sense RNA viruses reveal the complexity and dynamics of the cellular and viral epitranscriptomes during infection. Nucleic Acids Res 46:5776–5791.

23. Savidis G, McDougall WM, Meraner P, Perreira JM, Portmann JM, Trincucci G, John SP, Aker AM, Renzette N, Robbins DR, Guo Z, Green S, Kowalik TF, Brass AL. 2016. Identification of Zika Virus and Dengue Virus Dependency Factors using Functional Genomics. Cell Rep 16:232–246.

24. Hall JB, Allen FW. 1960. Studies on the incorporation of orotic acid into the 5-ribosyluracil phosphate of the ribonucleic acids of yeast. BBA - Biochimica et Biophysica Acta 39:557–559.

25. Nachtergaele S, He C. 2018. Chemical Modifications in the Life of an mRNA Transcript. Annu Rev Genet 52:349.

26. Kadumuri RV, Janga SC. 2018. Epitranscriptomic Code and Its Alterations in Human Disease. Trends Mol Med 24:886–903.

27. Delaunay S, Frye M. 2019. RNA modifications regulating cell fate in cancer. Nat Cell Biol 21:552–559.

28. Delaunay S, Helm M, Frye M. 2023. RNA modifications in physiology and disease: towards clinical applications. Nature Reviews Genetics 2023 25:104–122.

29. Roundtree IA, Evans ME, Pan T, He C. 2017. Dynamic RNA Modifications in Gene Expression Regulation. Cell 169:1187–1200.

30. Nachtergaele S, He C. 2016. The emerging biology of RNA post-transcriptional modifications. RNA Biol 14:156.

31. Zhang F, Kang Y, Wang M, Li Y, Xu T, Yang W, Song H, Wu H, Shu Q, Jin P. 2018. Fragile X mental retardation protein modulates the stability of its m6A-marked messenger RNA targets. Hum Mol Genet 27:3936–3950.

32. Roost C, Lynch SR, Batista PJ, Qu K, Chang HY, Kool ET. 2015. Structure and thermodynamics of N6-methyladenosine in RNA: A spring-loaded base modification. J Am Chem Soc 137:2107–2115.

33. Yustis JC, Devoucoux M, Côté J. 2024. The Functional Relationship Between RNA Splicing and the Chromatin Landscape. J Mol Biol 436:168614.

34. Kasowitz SD, Ma J, Anderson SJ, Leu NA, Xu Y, Gregory BD, Schultz RM, Wang PJ. 2018. Nuclear m6A reader YTHDC1 regulates alternative polyadenylation and splicing during mouse oocyte development. PLoS Genet 14:e1007412.

35. Amort T, Rieder D, Wille A, Khokhlova-Cubberley D, Riml C, Trixl L, Jia XY, Micura R, Lusser A. 2017. Distinct 5-methylcytosine profiles in poly(A) RNA from mouse embryonic stem cells and brain. Genome Biol 18:1.

36. Zheng G, Dahl JA, Niu Y, Fedorcsak P, Huang CM, Li CJ, Vågbø CB, Shi Y, Wang WL, Song SH, Lu Z, Bosmans RPG, Dai Q, Hao YJ, Yang X, Zhao WM, Tong WM, Wang XJ, Bogdan F, Furu K, Fu Y, Jia G, Zhao X, Liu J, Krokan HE, Klungland A, Yang YG, He C. 2013. ALKBH5 Is a Mammalian RNA Demethylase that Impacts RNA Metabolism and Mouse Fertility. Mol Cell 49:18–29.

37. Yang Y, Hsu PJ, Chen YS, Yang YG. 2018. Dynamic transcriptomic m6A decoration: Writers, erasers, readers and functions in RNA metabolism. Cell Res 28:616–624.

38. Lin S, Choe J, Du P, Triboulet R, Gregory RI. 2016. The m6A Methyltransferase METTL3 Promotes Translation in Human Cancer Cells. Mol Cell 62:335–345.

39. Boo SH, Kim YK. 2020. The emerging role of RNA modifications in the regulation of mRNA stability. Experimental & Molecular Medicine 52:400–408.

40. Shelton SB, Reinsborough C, Xhemalce B. 2016. Who Watches the Watchmen: Roles of RNA Modifications in the RNA Interference Pathway. PLoS Genet 12:e1006139.

41. Cappannini A, Ray A, Purta E, Mukherjee S, Boccaletto P, Moafinejad SN, Lechner A, Barchet C, Klaholz BP, Stefaniak F, Bujnicki JM. 2024. MODOMICS:ã database of RNA modificationsãnd related inf ormation. 2023 update. Nucleic Acids Res 52:D239–D244.

42. Arango D, Sturgill D, Alhusaini N, Dillman AA, Sweet TJ, Hanson G, Hosogane M, Sinclair WR, Nanan KK, Mandler MD, Fox SD, Zengeya TT, Andresson T, Meier JL, Coller J, Oberdoerffer S. 2018. Acetylation of Cytidine in mRNA Promotes Translation Efficiency. Cell 175:1872–1886.e24.

43. Bortolin-Cavaillé ML, Quillien A, Gamage ST, Thomas JM, Sas-Chen A, Sharma S, Plisson-Chastang C, Vandel L, Blader P, Lafontaine DLJ, Schwartz S, Meier JL, Cavaillé J. 2022. Probing small ribosomal subunit RNA helix 45 acetylation across eukaryotic evolution. Nucleic Acids Res 50:6284–6299.

44. Suzuki T, Ito S, Horikawa S, Suzuki T, Kawauchi H, Tanaka Y, Suzuki T. 2014. Human NAT10 is an ATP-dependent rna acetyltransferase responsible for N4-acetylcytidine formation in 18 S ribosomal RNA (rRNA). Journal of Biological Chemistry 289:35724–35730.

45. Thomas G, Gordon J, Rogg H. 1978. N4-Acetylcytidine. A previously unidentified labile component of the small subunit of eukaryotic ribosomes. Journal of Biological Chemistry 253:1101–1105.

46. Sharma S, Langhendries JL, Watzinger P, Kotter P, Entian KD, Lafontaine DLJ. 2015. Yeast Kre33 and human NAT10 are conserved 18S rRNA cytosine acetyltransferases that modify tRNAs assisted by the adaptor Tan1/THUMPD1. Nucleic Acids Res 43:2242–2258.

47. Tsai K, Jaguva Vasudevan AA, Martinez Campos C, Emery A, Swanstrom R, Cullen BR. 2020. Acetylation of Cytidine Residues Boosts HIV-1 Gene Expression by Increasing Viral RNA Stability. Cell Host Microbe 28:306–312.e6.

48. Hao H, Liu W, Miao Y, Ma L, Yu B, Liu L, Yang C, Zhang K, Chen Z, Yang J, Zheng Z, Zhang B, Deng F, Gong P, Yuan J, Hu Z, Guan W. 2022. N4-acetylcytidine regulates the replication and pathogenicity of enterovirus 71. Nucleic Acids Res 50:9339–9354.

49. Furuse Y. 2021. RNA modifications in genomic RNA of influenza a virus and the relationship between RNA modifications and viral infection. Int J Mol Sci 22:9127.

50. Gokhale NS, McIntyre ABR, McFadden MJ, Roder AE, Kennedy EM, Gandara JA, Hopcraft SE, Quicke KM, Vazquez C, Willer J, Ilkayeva OR, Law BA, Holley CL, Garcia-Blanco MA, Evans MJ, Suthar MS, Bradrick SS, Mason CE, Horner SM. 2016. N6-Methyladenosine in Flaviviridae Viral RNA Genomes Regulates Infection. Cell Host Microbe 20:654–665.

51. Lichinchi G, Zhao BS, Wu Y, Lu Z, Qin Y, He C, Rana TM. 2016. Dynamics of Human and Viral RNA Methylation during Zika Virus Infection. Cell Host Microbe 20:666–673.

52. Li Z, Lee I, Moradi E, Hung NJ, Johnson AW, Marcotte EM. 2009. Rational Extension of the Ribosome Biogenesis Pathway Using Network-Guided Genetics. PLoS Biol 7:e1000213.

53. Münster M, Płaszczyca A, Cortese M, Neufeldt CJ, Goellner S, Long G, Bartenschlager R. 2018. A Reverse Genetics System for Zika Virus Based on a Simple Molecular Cloning Strategy. Viruses 10:368.

54. Donald CL, Brennan B, Cumberworth SL, Rezelj V V., Clark JJ, Cordeiro MT, Freitas de Oliveira França R, Pena LJ, Wilkie GS, Da Silva Filipe A, Davis C, Hughes J, Varjak M, Selinger M, Zuvanov L, Owsianka AM, Patel AH, McLauchlan J, Lindenbach BD, Fall G, Sall AA, Biek R, Rehwinkel J, Schnettler E, Kohl A. 2016. Full Genome Sequence and sfRNA Interferon Antagonist Activity of Zika Virus from Recife, Brazil. PLoS Negl Trop Dis 10:e0005048.

55. Mufrrih M, Chen B, Chan S-W. 2021. Zika Virus Induces an Atypical Tripartite Unfolded Protein Response with Sustained Sensor and Transient Effector Activation and a Blunted BiP Response. mSphere 6:e0036121.

56. Frumence E, Roche M, Krejbich-Trotot P, El-Kalamouni C, Nativel B, Rondeau P, Missé D, Gadea G, Viranaicken W, Desprès P. 2016. The South Pacific epidemic strain of Zika virus replicates efficiently in human epithelial A549 cells leading to IFN-β production and apoptosis induction. Virology 493:217–226.

57. Hou W, Armstrong N, Obwolo LA, Thomas M, Pang X, Jones KS, Tang Q. 2017. Determination of the Cell Permissiveness Spectrum, Mode of RNA Replication, and RNA-Protein Interaction of Zika Virus. BMC Infect Dis 17:239.

58. Li C, Deng YQ, Wang S, Ma F, Aliyari R, Huang XY, Zhang NN, Watanabe M, Dong HL, Liu P, Li XF, Ye Q, Tian M, Hong S, Fan J, Zhao H, Li L, Vishlaghi N, Buth JE, Au C, Liu Y, Lu N, Du P, Qin FXF, Zhang B, Gong D, Dai X, Sun R, Novitch BG, Xu Z, Qin CF, Cheng G. 2017. 25-Hydroxycholesterol Protects Host against Zika Virus Infection and Its Associated Microcephaly in a Mouse Model. Immunity 46:446–456.

59. Berglund G, Lennon CD, Badu P, Berglund JA, Pager CT. 2024. Transcriptomic Signatures of Zika Virus Infection in Patients and a Cell Culture Model. Microorganisms 12:1499.

60. Mesa RA, Yasothan U, Kirkpatrick P. 2012. Ruxolitinib. Nat Rev Drug Discov 11:103–104.

61. Otter CJ, Bracci N, Parenti NA, Ye C, Asthana A, Blomqvist EK, Tan LH, Pfannenstiel JJ, Jackson N, Fehr AR, Silverman RH, Burke JM, Cohen NA, Martinez-Sobrido L, Weiss SR. 2024. SARS-CoV-2 nsp15 endoribonuclease antagonizes dsRNA-induced antiviral signaling. Proc Natl Acad Sci U S A 121:e2320194121.

62. Dang Y, Li J, Li Y, Wang Y, Zhao Y, Zhao N, Li W, Zhang H, Ye C, Ma H, Zhang L, Liu H, Dong Y, Yao M, Lei Y, Xu Z, Zhang F, Ye W. 2024. N-acetyltransferase 10 regulates alphavirus replication via N4-acetylcytidine (ac4C) modification of the lymphocyte antigen six family member E (LY6E) mRNA. J Virol 98:e0135023.

63. Li P, Jiang H, Peng H, Zeng W, Zhong Y, He M, Xie L, Chen J, Guo D, Wu J, Li CM. 2021. Non-structural protein 5 of Zika virus interacts with p53 in human neural progenitor cells and induces p53-mediated apoptosis. Virol Sin 36:1411–1420.

64. Teng Y, Liu S, Guo X, Liu S, Jin Y, He T, Bi D, Zhang P, Lin B, An X, Feng D, Mi Z, Tong Y. 2017. An integrative analysis reveals a central role of P53 activation via MDM2 in Zika virus infection induced cell death. Front Cell Infect Microbiol 7:327.

65. El Ghouzzi V, Bianchi FT, Molineris I, Mounce BC, Berto GE, Rak M, Lebon S, Aubry L, Tocco C, Gai M, Chiotto AMA, Sgrò F, Pallavicini G, Simon-Loriere E, Passemard S, Vignuzzi M, Gressens P, Di Cunto F. 2016. ZIKA virus elicits P53 activation and genotoxic stress in human neural progenitors similar to mutations involved in severe forms of genetic microcephaly and p53. Cell Death & Disease 7:e2440–e2440.

66. Zhang Y, Lu L, Li X. 2022. Detection technologies for RNA modifications. Experimental & Molecular Medicine 54:1601–1616.

67. Arango D, Sturgill D, Yang R, Kanai T, Bauer P, Roy J, Wang Z, Hosogane M, Schiffers S, Oberdoerffer S. 2022. Direct epitranscriptomic regulation of mammalian translation initiation through N4-acetylcytidine. Mol Cell 82:2797–2814.e11.

68. Zhang X, Qin S, Huang F, Liu H, Wang J, Chen Z, Hao H, Ding S, Liu L, Yu B, Liu Y, Liu H, Guan W. 2025. N4-acetylcytidine coordinates with NP1 and CPSF5 to facilitate alternative RNA processing during the replication of minute virus of canines. Nucleic Acids Res 53:gkaf229.

69. Yan Q, Zhou J, Gu Y, Huang W, Ruan M, Zhang H, Wang T, Wei P, Chen G, Li W, Lu C. 2024. Lactylation of NAT10 promotes N4-acetylcytidine modification on tRNASer-CGA-1-1 to boost oncogenic DNA virus KSHV reactivation. Cell Death Differ 31:1362–1374.

70. Zhang S, Huang F, Wang Y, Long Y, Li Y, Kang Y, Gao W, Zhang X, Wen Y, Wang Y, Pan L, Xia Y, Yang Z, Yang Y, Mo H, Li B, Hu J, Song Y, Zhang S, Dong S, Du X, Li Y, Liu Y, Liao W, Gao Y, Zhang Y, Chen H, Liang Y, Chen J, Weng H, Huang H. 2024. NAT10-mediated mRNA N4-acetylcytidine reprograms serine metabolism to drive leukaemogenesis and stemness in acute myeloid leukaemia. Nature Cell Biology 26:2168–2182.

71. Wilhelm E, Poirier M, Rocha M Da, Bédard M, McDonald PP, Lavigne P, Hunter CL, Bell B. 2024. Mitotic deacetylase complex (MiDAC) recognizes the HIV-1 core promoter to control activated viral gene expression. PLoS Pathog 20:e1011821.

72. Sun L, Li X, Xu F, Chen Y, Li X, Yang Z, Yang Y, Wang K, Ren T, Lin Z, Wang H, Wang X, Lu Y, Song Z, Cheng ZL, Wu D. 2025. A critical role of N4-acetylation of cytidine in mRNA by NAT10 in T cell expansion and antiviral immunity. Nat Immunol 26:619–634.

73. Yan Q, Zhou J, Wang Z, Ding X, Ma X, Li W, Jia X, Gao SJ, Lu C. 2023. NAT10-dependent N 4-acetylcytidine modification mediates PAN RNA stability, KSHV reactivation, and IFI16-related inflammasome activation. Nat Commun 14:6327.

74. Shah PS, Link N, Jang GM, Sharp PP, Zhu T, Swaney DL, Johnson JR, Von Dollen J, Ramage HR, Satkamp L, Newton B, Hüttenhain R, Petit MJ, Baum T, Everitt A, Laufman O, Tassetto M, Shales M, Stevenson E, Iglesias GN, Shokat L, Tripathi S, Balasubramaniam V, Webb LG, Aguirre S, Willsey AJ, Garcia-Sastre A, Pollard KS, Cherry S, Gamarnik A V., Marazzi I, Taunton J, Fernandez-Sesma A, Bellen HJ, Andino R, Krogan NJ. 2018. Comparative Flavivirus-Host Protein Interaction Mapping Reveals Mechanisms of Dengue and Zika Virus Pathogenesis. Cell 175:1931–1945.e18.

75. Zhang H, Chen Z, Zhou J, Gu J, Wu H, Jiang Y, Gao S, Liao Y, Shen R, Miao C, Chen W. 2022. NAT10 regulates neutrophil pyroptosis in sepsis via acetylating ULK1 RNA and activating STING pathway. Communications Biology 5:1–13.

76. Liu WC, Wei YH, Chen JF, Xing XL, Jia HX, Yang XY, Huang YJ, Liu XM, Xiao K, Guo XD, Can C, Zhang AM, He N, Zhang HL, Ma DX. 2025. Inhibition of tumor-intrinsic NAT10 enhances antitumor immunity by triggering type I interferon response via MYC/CDK2/DNMT1 pathway. Nature Communications 16:1–21.

77. Serman T, Chiang C, Liu GQ, Sayyad Z, Pandey S, Volcic M, Lee H, Muppala S, Acharya D, Goins C, Stauffer SR, Sparrer KMJ, Gack MU. 2023. Acetylation of the NS3 helicase by KAT5γ is essential for flavivirus replication. Cell Host Microbe 31:1317–1330.e10.

78. Fagre AC, Lewis J, Miller MR, Mossel EC, Lutwama JJ, Nyakarahuka L, Nakayiki T, Kityo R, Nalikka B, Towner JS, Amman BR, Sealy TK, Foy B, Schountz T, Anderson J, Kading RC. 2021. Subgenomic flavivirus RNA (sfRNA) associated with Asian lineage Zika virus identified in three species of Ugandan bats (family Pteropodidae). Scientific Reports 11:1–8.

79. Graham ME, Merrick C, Akiyama BM, Szucs MJ, Leach S, Kieft JS, Beckham JD. 2023. Zika virus dumbbell-1 structure is critical for sfRNA presence and cytopathic effect during infection. mBio 14:e0110823.

80. Schwarz MC, Sourisseau M, Espino MM, Gray ES, Chambers MT, Tortorella D, Evans MJ. 2016. Rescue of the 1947 Zika virus prototype strain with a cytomegalovirus promoter-driven cDNA clone. mSphere 1:e00246–16.

81. Ratnadiwakara M, Änkö M-L. 2018. mRNA Stability assay using transcription inhibition by actinomycin D in mouse pluripotent stem cells. Bio Protoc 8:e3072.

82. Biegel JM, Henderson E, Cox EM, Bonenfant G, Netzband R, Kahn S, Eager R, Pager CT. 2017. Cellular DEAD-box RNA helicase DDX6 modulates interaction of miR-122 with the 5′ untranslated region of hepatitis C virus RNA. Virology 507:231–241.

83. Sas-Chen A, Thomas JM, Matzov D, Taoka M, Nance KD, Nir R, Bryson KM, Shachar R, Liman GLS, Burkhart BW, Gamage ST, Nobe Y, Briney CA, Levy MJ, Fuchs RT, Robb GB, Hartmann J, Sharma S, Lin Q, Florens L, Washburn MP, Isobe T, Santangelo TJ, Shalev-Benami M, Meier JL, Schwartz S. 2020. Dynamic RNA acetylation revealed by quantitative cross-evolutionary mapping. Nature 583:638–643.

